# Poxvirus-encoded TNF receptor homolog dampens inflammation and protects the host from uncontrolled lung pathology and death during respiratory infection

**DOI:** 10.1101/2020.02.24.963520

**Authors:** Zahrah Al Rumaih, Ma. Junaliah Tuazon Kels, Esther Ng, Pratikshya Pandey, Sergio M. Pontejo, Alí Alejo, Antonio Alcamí, Geeta Chaudhri, Gunasegaran Karupiah

## Abstract

Ectromelia virus (ECTV) causes mousepox, a surrogate mouse model for smallpox caused by variola virus in humans. Both viruses encode tumor necrosis factor receptor (TNFR) homologs termed cytokine response modifier (Crm) proteins, containing a TNF-binding domain and a chemokine-binding domain termed smallpox virus-encoded chemokine receptor (SECRET) domain. Infection of ECTV-resistant C57BL/6 mice with an ECTV CrmD deletion mutant resulted in uniform mortality due to excessive TNF secretion and dysregulated inflammatory cytokine production but viral load was not affected. CrmD dampened lung pathology, leukocyte recruitment and inflammatory cytokines including TNF, IL-6, IL-10 and IFN-γ. Blockade of IL-6, IL-10R or TNF function with monoclonal antibodies reduced lung pathology and provided 60-100% protection from an otherwise lethal infection. IFN-γ caused lung pathology only when both the TNF-binding and SECRET domains were deleted but it was neither necessary nor sufficient to cause pathology when only the CrmD SECRET domain was expressed by virus.

## Introduction

Tumor necrosis factor (TNF) plays key roles in the maintenance of homeostasis, acute inflammation (Arnett et al., 2001; Korner et al., 1997; Marino et al., 1997; Papathanasiou et al., 2015; Pasparakis et al., 1996), and anti-microbial defense (Bean et al., 1999; Domm et al., 2008; Kalliolias and Ivashkiv, 2016; Wilhelm et al., 2001). This cytokine can also cause serious pathology during acute viral infections and chronic inflammatory diseases (Belisle et al., 2010; Szretter et al., 2007; Tracey et al., 1987). TNF is produced early during the course of a viral infection following activation of the nuclear factor-kappa B (NF-κB) inflammatory pathway. It is expressed as a transmembrane protein (mTNF), which is cleaved by metalloproteinase enzymes to produce soluble TNF (sTNF) (Black et al., 1997; Moss et al., 1997). The two forms of TNF play important roles in the inflammatory response to viral infections and exert their activities by signaling through TNF receptor I (TNFRI) or TNFRII (Beutler and Cerami, 1989; Chu, 2013). Whereas both forms of TNF are capable of activating the TNFRI signaling pathway, TNFRII is activated more efficiently by mTNF (Grell et al., 1995). Furthermore, mTNF can also participate in reverse signaling, a process whereby the interaction of mTNF with TNFR leads to the transmission of signals in the direction of the cell expressing mTNF (Horiuchi et al., 2010; Juhasz et al., 2013; Watts et al., 1999).

Many viruses, including orthopoxviruses (OPV), encode proteins that can modulate the host TNF response (Alcami, 2003; Alcami and Koszinowski, 2000; Alejo et al., 2018; Seet et al., 2003; Stanford et al., 2007). OPV-encoded homologs of mammalian TNF receptor (TNFR) can modulate inflammation and the immune response (Alejo et al., 2018), a strategy that provides an advantage to the virus, allowing successful replication and transmission to other hosts. There are four types of viral TNFR (vTNFR), also known as cytokine response modifier B (CrmB), CrmC, CrmD and CrmE (Pontejo et al., 2015a; Smith, 1996). CrmC and CrmE have specificity for TNF (Reading et al., 2002; Saraiva and Alcami, 2001; Smith et al., 1996), whereas CrmB and CrmD bind TNF (Hu et al., 1994; Loparev et al., 1998) as well as lymphotoxin (LT)-α and -β (Alejo et al., 2006; Pontejo et al., 2015a). All Crm proteins contain a TNF-binding domain at the N-terminal, consisting of four cysteine-rich domains (CRDs), which are homologous to mammalian TNFR CRD (Lefkowitz et al., 2005; Ribas et al., 2003) and shown to bind to mTNF and inhibit its ability to activate host TNFRs (Pontejo et al., 2015b). In addition, CrmB, encoded by variola virus (VARV, agent of smallpox), and CrmD, encoded by ectromelia virus (ECTV; agent of mousepox) contain an extended C-terminal chemokine-binding domain known as the smallpox virus-encoded chemokine receptor (SECRET) domain (Alejo et al., 2006).

ECTV is a strict mouse pathogen and the disease it causes in mice is very similar to smallpox caused by VARV in humans. ECTV and VARV are closely related OPV, and mousepox has been utilized extensively as a small animal model to understand, among others, the pathogenetic basis for resistance and susceptibility of humans to smallpox. Among inbred strains of mice, resistance to mousepox requires robust inflammatory and polarized T helper 1-like immune responses, which are associated with the cytokines interleukin 2 (IL-2), interferon gamma (IFN-γ) and TNF, and robust natural killer (NK) cell, cytotoxic T lymphocyte (CTL) and neutralizing antibody responses (Chaudhri et al., 2006; Chaudhri et al., 2004; Fang and Sigal, 2005; Karupiah et al., 1996; Parker et al., 2007). The C57BL/6 strain is resistant to ECTV infection but deficiency in IFN (Chaudhri et al., 2004; Karupiah et al., 1993; Panchanathan et al., 2005) or TNF (Kels et al., 2019) makes it highly susceptible to infection. ECTV-susceptible mice such as the BALB/c strain generate weak inflammatory and immune responses, associated with sub-optimal IFN-γ, TNF, NK cell and CTL responses. However, infection of this strain with a CrmD deletion mutant virus (ECTV^ΔCrmD^) augments inflammation, NK cell and CTL activities, resulting in effective control of virus replication and survival (Alejo et al., 2018). CrmD clearly provides an advantage to ECTV in the BALB/c strain.

In this report, we have investigated the consequences of infecting the ECTV-resistant C57BL/6 mouse strain with ECTV^ΔCrmD^ or a virus that expresses only the SECRET domain (ECTV^Rev.SECRET^). As the TNF-binding domain of CrmD can also interact with LT-α and LT-β, we also determined the outcomes of infecting mice deficient in TNF production or those that only express mTNF. We show here that CrmD plays an essential role in dampening inflammation, involving reverse signaling through mTNF in C57BL/6 mice during a respiratory infection. Deficiency in CrmD substantially increased the susceptibility of C57BL/6 mice without any increases in viral load but significantly augmented levels of several cytokines including TNF, IL-6, IL-10 and IFN-γ. The increased levels of these cytokines were responsible for lung pathology as blockade of TNF, IL-6 or IL-10 with monoclonal antibodies (mAb) resulted in reduced inflammation, improved lung pathology and 60-100% survival of mice infected with either ECTV^ΔCrmD^ or ECTV^Rev.SECRET^. A requirement for IFN-γ in causing lung pathology was dependent on the type of virus used for infection. In ECTV^ΔCrmD^-infected C57BL/6 mice, neutralization of IFN-γ significantly reduced lung pathology and viral load with 100% survival from an otherwise lethal infection whereas anti-IFN-γ treatment had minimal effects in ECTV^Rev.SECRET^-infected C57BL/6 mice. Thus, IFN-γ causes pathology when both the TNF-binding and SECRET domains are deleted whereas it is insufficient to cause pathology when only the SECRET domain is present. The recovery or death from ECTV infection is determined not only by a balance between the host’s ability to produce TNF and the virus’ ability to neutralize its effects but also how CrmD signaling through mTNF affects production of other inflammatory cytokines.

## Results

### Absence of CrmD increases susceptibility of C57BL/6 mice to infection with ECTV

Mutant viruses lacking both copies of CrmD (ECTV^ΔCrmD^) or expressing only the SECRET molecule (ECTV^Rev.SECRET^), and wild-type ECTV (ECTV^WT^) were used to study the role of CrmD in the pathogenesis of mousepox in ECTV-resistant C57BL/6 mice.

We used wild-type (WT) mice, TNF deficient (TNF^-/-^) mice and triple mutant (TM) mice, which were engineered to express only non-cleavable mTNF but not TNFRI or TNFRII (mTNF^Δ/Δ^.TNFRI^-/-^.TNFRII^-/-^) mice. TNF^-/-^ and TM mice were used to determine whether the presence or absence of CrmD or the SECRET domain impacted on the pathogenesis of ECTV infection in the absence of TNF or in the presence of only mTNF, respectively. The TM mice are similar to TNF^-/-^ mice in so far as they are devoid of an endogenous TNF response but unlike the latter, TM mice can respond to exogenous TNFR or CrmD. The TM mice are marginally more susceptible than mTNF^Δ/Δ^ mice, which express both TNFRI and TNFRII, to ECTV^WT^ infection (Figure S1). We predicted that CrmD binding to sTNF in WT mice can reduce the bioavailability of the cytokine whereas its binding to mTNF can potentially modulate the inflammatory response further through reverse signaling (Horiuchi et al., 2010; Juhasz et al., 2013; Watts et al., 1999) in ECTV^WT^-infected WT and TM mice but not in TNF^-/-^ mice.

Groups of mice were inoculated intranasally (i.n.) with ECTV^WT^, ECTV^ΔCrmD^ or ECTV^Rev.SECRET^ and monitored daily for changes in body weight and disease signs, presented as clinical scores using a scoring system (Table S1).

WT mice infected with the mutant viruses exhibited significant weight loss and higher clinical scores compared with ECTV^WT^-infected animals from day 7 post-infection (p.i.) (Figures 1A and 1B). Clinical scores between groups of WT mice infected with either of the mutant viruses were comparable. However, ECTV^Rev.SECRET^-infected mice exhibited significantly higher weight loss at days 7-9 compared to ECTV^WT^-infected mice. TNF^-/-^ mice fared poorly regardless of the type of virus used for infection and the presence or absence of CrmD or SECRET did not influence weight loss or clinical scores (Figures 1C and 1D). TM mice infected with ECTV^WT^ lost less weight than those infected with mutant viruses at days 8 and 9 but they were not statistically significant (Figure 1E). The clinical scores in TM mice infected with any of the viruses were also similar until day 8 but by day 9, the clinical scores of ECTV^WT^-infected mice were significantly lower than ECTV^ΔCrmD^-infected animals (Figures 1E and 1F). WT and TM mice infected with ECTV^WT^ fared much better than all 3 strains infected with mutant viruses. Although no mortality was recorded in this experiment, all animals were euthanized at day 9 p.i. due to ethical and animal welfare reasons.

**Figure 1.**
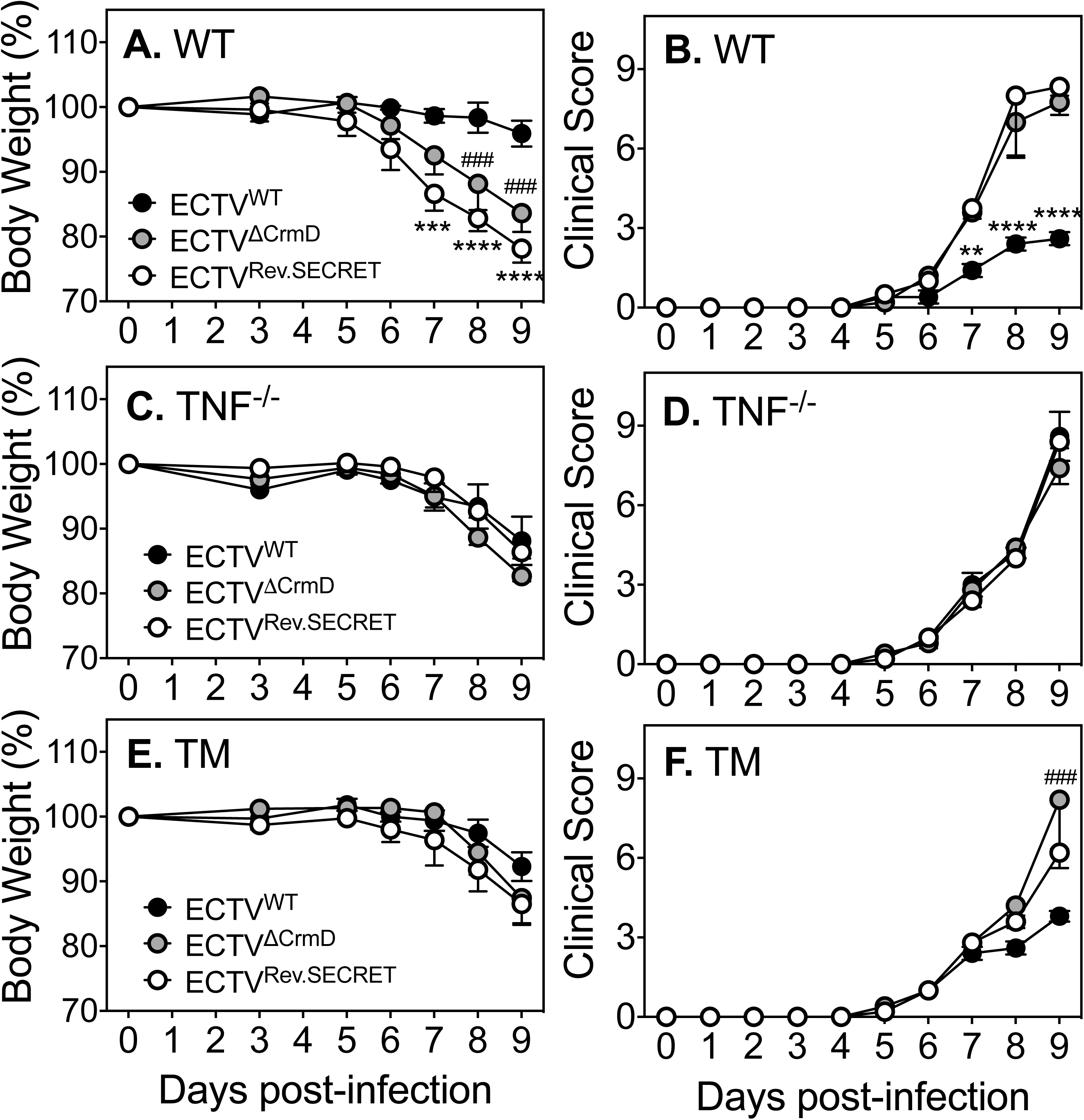
Increased susceptibility of WT and TM mice to ECTV^ΔCrmD^ and ECTV^Rev.SECRET^. Groups of WT, TNF^-/-^ and TM female mice (5 mice/group) were infected i.n. with 25 plaque forming units (PFU) of ECTV^WT^, ECTV^ΔCrmD^ or ECTV^Rev.SECRET^. Weights (A, C and E) and clinical scores (B, D and F) were monitored during the course of infection. Mice were sacrificed at day 9 p.i. Data for weights and clinical scores are expressed as means ± SEM. Statistical analysis was done using two way-ANOVA followed by Holm-Sidak’s post-tests. Weight loss and clinical scores in mice infected with ECTV^WT^ was compared to mice infected with the mutant viruses. *, p < 0.05; **, p < 0.01; ***, p < 0.001; ^###^, p < 0.001; ****, p < 0.0001. Data shown are from one of 3 separate experiments with comparable outcomes. See also Table S1.

The results indicated that CrmD plays an important role in reducing weight loss and clinical symptoms during respiratory ECTV infection. Infection of WT and TM animals with CrmD deletion mutant ECTV had a significant impact on the overall heath deterioration contributing to significant morbidity. The data also confirmed that TNF is critical for CrmD to manifest its biological effects.

### CrmD deficiency exacerbates lung pathology but does not affect viral load in C57BL/6 mice

We investigated whether the significant morbidity among WT and TM animals infected with mutant viruses was due to exacerbated lung pathology, increased viral load or both. Mice were infected i.n. with ECTV^WT^, ECTV^ΔCrmD^ or ECTV^Rev.SECRET^, sacrificed at day 9 p.i. and lung tissue was collected for histology and measurement of viral load.

WT mice infected with ECTV^WT^ exhibited minimal parenchymal and perivascular edema with minor damage to the epithelial lining of the bronchioles, and minimal diffusion of cellular infiltrates as revealed by hematoxylin and eosin (H&E)-stained lung sections (Figure 2A). In contrast, ECTV^WT^-infected TNF^-/-^ mice exhibited severe lung injury with total destruction of most alveoli due to fluid accumulation, diffuse to multifocal infiltration of leukocytes and extensive damage to the entire epithelial lining of the bronchioles (Figure 2B). Lungs from TM mice infected with ECTV^WT^ exhibited more severe pathology than WT lungs but somewhat less severe than the pathology in TNF^-/-^ mice (Figure 2C).

**Figure 2.**
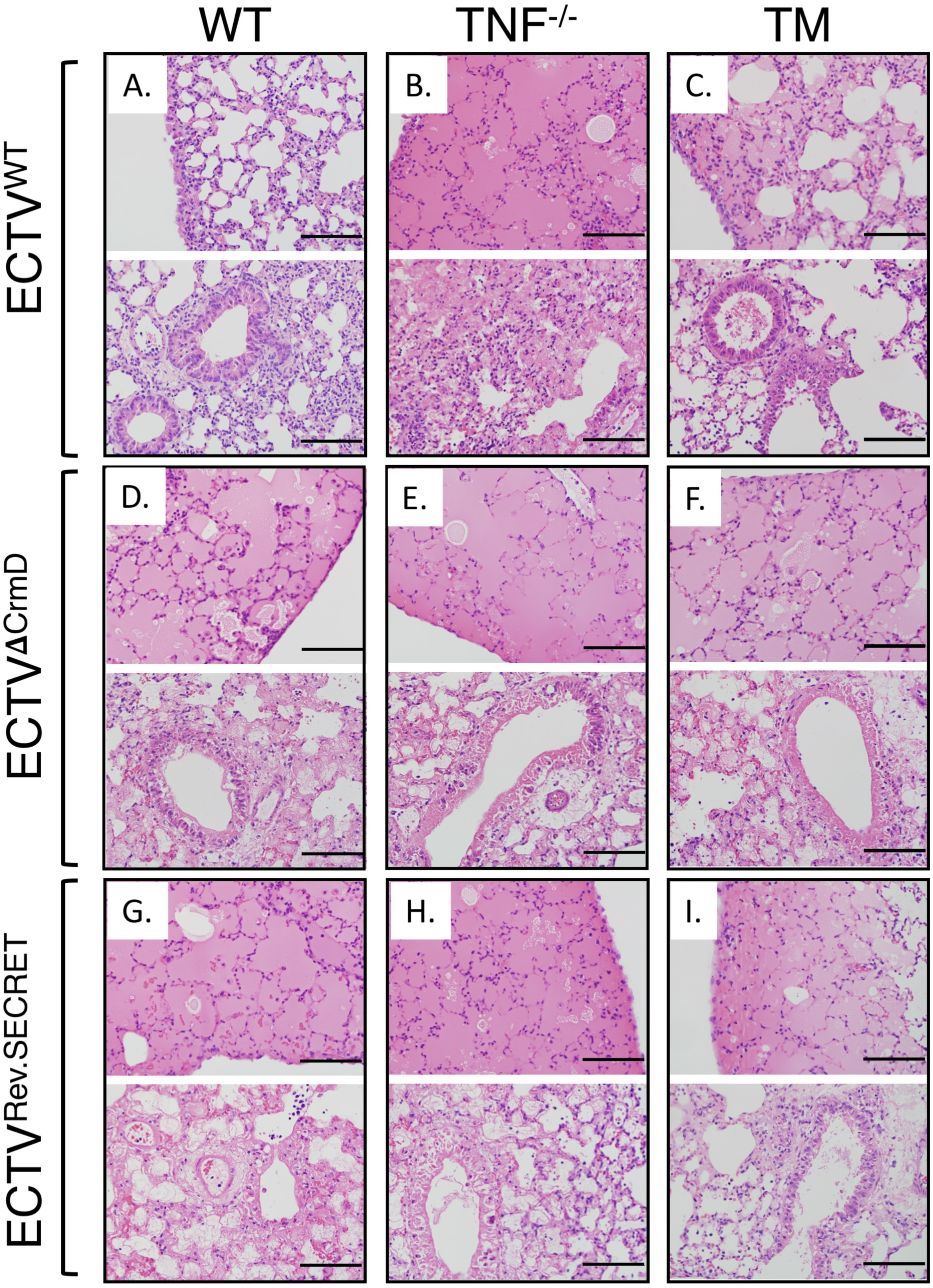
Exacerbated lung pathology in mice infected with ECTV^ΔCrmD^ or ECTV^Rev.SECRET^. Groups of WT, TNF^-/-^ and TM female mice (5 mice/group) were infected i.n. with 25 PFU of ECTV^WT^, ECTV^ΔCrmD^ or ECTV^Rev.SECRET^. Representative lung histology sections from WT (A, D, G), TNF^-/-^ (B, E, H) and TM (C, F, I) mice at 9 days p.i. Slides were examined on all fields at 400x magnification. Representative sections are from one of 3 individual experiments with comparable outcomes. Bars correspond to 100 µm.

Infection with ECTV^ΔCrmD^ exacerbated the lung pathology in WT mice with massive fluid accumulation, destruction of most alveoli, minimal diffusion to multifocal infiltration of leukocytes and significant damage to the epithelial lining of the bronchioles (Figure 2D). The absence of CrmD did not impact on lung pathology in TNF^-/-^ mice (Figure 2E). Lung pathology in TM mice infected with ECTV^ΔCrmD^ was similar to the pathology caused by the mutant virus in WT mice (Figure 2F).

The lung pathology caused by ECTV^Rev.SECRET^ (Figures 2G-2I) was similar to the pathology caused by infection with ECTV^ΔCrmD^ (Figures 2D-2F) in all three strains of mice. Most of the bronchial walls exhibited either complete or nearly complete epithelial necrosis, there were interstitial infiltration of leukocytes and alveolar septal wall thickening in WT (Figure 2G), TNF^-/-^ (Figure 2H) and TM lungs (Figure 2I). Moreover, fluid extravasated to large portions of the lobes, leading to the collapse of the alveolar septal walls in all strains of mice.

We semi-quantified the histopathological changes in the lungs using a scoring system based on the degree of edema, bronchial epithelial necrosis, parenchymal and perivascular inflammatory infiltrates and alveolar septal wall damage (Table S2). Five distinct observations were made from the analysis. First, lung sections from WT mice had the lowest scores, TM lung sections had intermediate scores and TNF^-/-^ lung sections had the highest scores in response to ECTV^WT^ infection (Figure 3A). Second, infection of WT mice with either mutant virus substantially increased lung histopathological scores. Third, in TNF^-/-^ mice, all three viruses caused considerable levels of lung damage with no significant differences in the histopathological scores. Fourth, TM lung sections from mice infected with ECTV^ΔCrmD^ or ECTV^Rev.SECRET^ had significant increases in their histopathological scores compared to those infected with ECTV^WT^. Finally, the mutant viruses caused comparable levels of lung damage in all three strains of mice.

**Figure 3.**
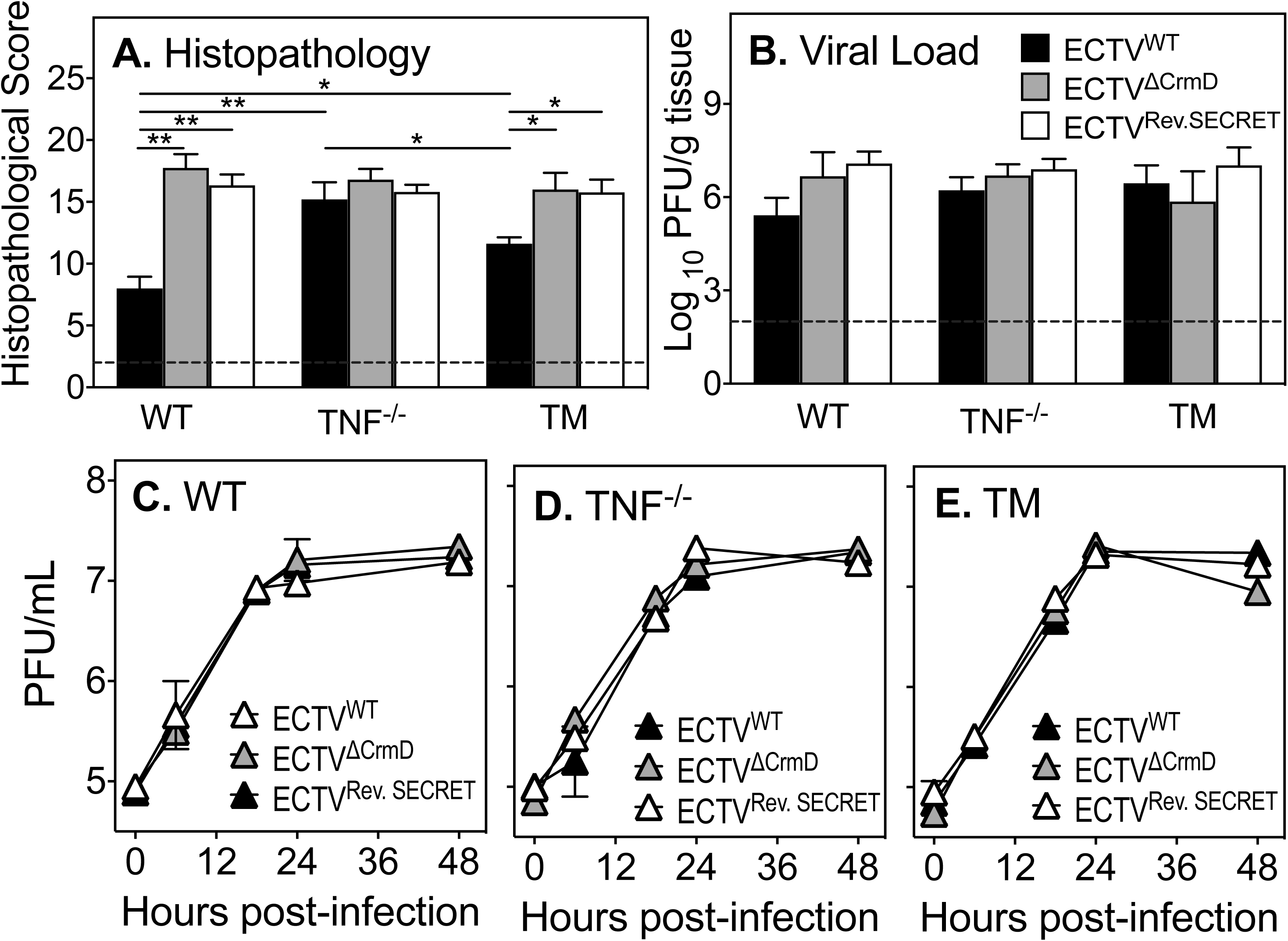
CrmD deficiency exacerbates lung pathology in WT and TM mice but has no effect on viral load. Groups of WT, TNF^-/-^ and TM mice (5 mice/group) were infected i.n. with 25 PFU ECTV^WT^, ECTV^ΔCrmD^ or ECTV^Rev.SECRET^. (A) Histopathological scores of lung sections from mice 9 days p.i. Slides were examined on all fields at 400x magnification. Data are expressed as means ± SEM and statistical analysis was done using ordinary one-way ANOVA test followed by Tukey’s multiple comparisons tests, where *, p < 0.05 and **, p < 0.01. (B). Lung viral load at day 9 p.i. Data was log transformed and expressed as means ± SEM. Statistical analysis was undertaken using ordinary one-way ANOVA followed by Fisher’s least significant difference tests. Data are from one of 2 separate experiments with comparable outcomes. Broken line corresponds to the limit of virus detection. *In vitro* virus replication in BMDM from (C) WT, (D) TNF^-/-^ and (E) TM mice. BMDM were infected at a multiplicity of infection of 1 with ECTV^WT^, ECTV^ΔCrmD^ or ECTV^Rev.SECRET^. Cells were harvested at 0, 6, 12, 24 and 48 h p.i. and viral load was determined by plaque assays. Data were log transformed and expressed as means ± SEM of duplicate samples for each time point. Statistical analysis was achieved using two-way ANOVA followed by Tukey’s post-tests. There were no significant differences in titers between the indicated viruses regardless of the genotype of BMDM used. Data shown are from one experiment.

The lung viral load in all three strains of mice infected with ECTV^WT^, ECTV^ΔCrmD^ or ECTV^Rev.SECRET^ were comparable as the titers of viruses within each strain or across strains were not statistically different. This finding indicated that the TNF-binding or SECRET domains had no effect on viral replication *in vivo* (Figure 3B). This result also indicated that the cause of severe lung pathology in mice infected with mutant viruses was not related to viral load. *In vitro*, the replication kinetics of ECTV^WT^, ECTV^ΔCrmD^ or ECTV^Rev.SECRET^ in bone marrow-derived macrophages (BMDM) from WT (Figure 3C), TNF^-/-^ (Figure 3D) or TM (Figure 3E) mice over a 48-h period were comparable, consistent with the *in vivo* results that indicated neither the TNF-binding domain nor the SECRET domain influence virus replication.

### The effect of CrmD deficiency on leukocyte recruitment/ infiltration

We used flow cytometry of collagenase-digested lungs from uninfected and virus-infected mice to ascertain whether the TNF-binding domain and/or the SECRET domain affected leukocyte recruitment into the lungs, which might have contributed to pathology. We enumerated the total number of leukocytes and leukocyte subsets at early (day 4) and later (day 8) stages of infection.

At day 4 p.i., numbers of leukocytes or leukocyte subsets did not increase significantly in WT animals following infection with ECTV^WT^ when compared with naïve animals (Figures 4A-K). However, in the lungs of ECTV^ΔCrmD^-infected animals, there were significant reductions in numbers of leukocytes (CD45.2^+^) (Figure 4A) and leukocyte subsets [T cells (CD3^+^) (Figure 4B), T cell subsets [(CD3^+^CD8^+^, CD3^+^CD4^+^, CD3^+^TCR γδ^+^) (Figures 4C-E), B cells (CD45R^+^) (Figure 4F), NK cells (NK1.1^+^) (Figure 4H), neutrophils (CD11b^hi^Ly6c^low^) (Figure 4J)] and inflammatory monocytes (CD11b^hi^Ly6c^hi^) (Figure 4K) compared to those infected with ECTV^WT^ or ECTV^Rev.SECRET^. Although the absence of CrmD did not affect numbers of macrophages (F4/80^+^) (Figure 4G) or dendritic cells (CD11c^+^) (Figure 4I), the presence of only the SECRET domain in ECTV^Rev.SECRET^-infected animals significantly increased numbers of these subsets. In fact, the SECRET domain appeared to be critical for leukocyte recruitment into the lungs of infected WT mice.

**Figure 4.**
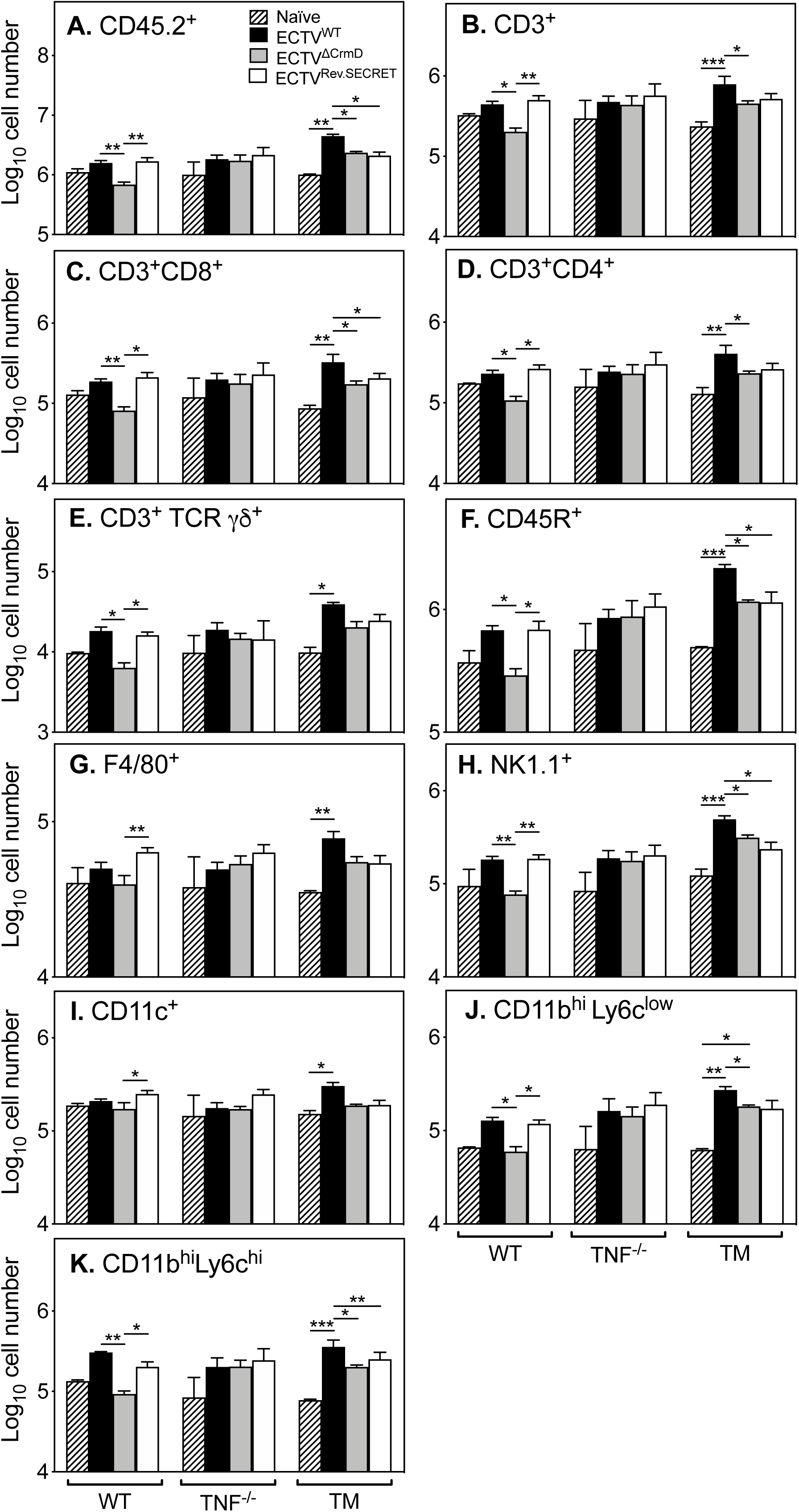
The SECRET domain of CrmD is important for leukocyte recruitment to the lungs early during the course of infection. Groups of WT, TNF^-/-^ and TM mice (5 mice/group) were infected i.n. with 25 PFU ECTV^WT^, ECTV^ΔCrmD^ or ECTV^Rev.SECRET^. At day 4 p.i., lungs were collected from virus-infected mice and 2 naïve animals from each group. Single cell suspensions of digested lungs were stained with fluorochrome-conjugated antibodies and analyzed by flow cytometry. Cell numbers were calculated based on yields per digested lung. Leukocytes were gated on CD45^+^ cells (CD45.2^+^). Shown are numbers of T cells (CD3^+^), T cell subsets (CD3^+^ CD4^+^, CD3^+^ CD8^+^ or CD3^+^ TCR γδ^+^), B cells (CD45R^+^), macrophages (F4/80^+^), NK cells (NK1.1^+^), dendritic cells (CD11c^+^), inflammatory monocytes (CD45^+^ CD11b^hi^ Ly6c^hi^) and neutrophils (CD45^+^ CD11b^hi^ Ly6c^low^). Data were log transformed and expressed as means ± SEM. Statistical significance was obtained using the two-way ANOVA followed by Tukey’s multiple comparisons tests, where *, p < 0.05; **, p < 0.01; ***, p < 0.001. Data shown are from one of 2 separate experiments with comparable results.

In contrast to WT mice, TNF^-/-^ animals did not exhibit any statistically significant increases or decreases in leukocyte subsets present in the lungs regardless of the type of infecting virus. This finding further indicated that TNF is critical for the function of CrmD and that any biological consequences of interaction of CrmD with LT-α or LT-β had a minimal impact on leukocyte recruitment to the lungs.

In TM mice, the total infiltrating leukocytes (CD45.2^+^) (Figure 4A) and most of the subsets examined were significantly higher in numbers in lungs of ECTV^WT^-infected animals compared to uninfected or mutant virus-infected animals. The absence of CrmD reduced leukocyte subset numbers in ECTV^ΔCrmD^-infected mice, similar to the outcome in WT mice. However, unlike in WT mice, the presence of only the SECRET domain in ECTV^Rev.SECRET^-infected mice did not contribute to any increases in numbers of leukocyte subsets.

At day 8 p.i., leukocyte numbers in WT lungs were comparable irrespective of the infecting virus (Figure S2). Similarly, in TNF^-/-^ mice, there were no differences in numbers and types of leukocytes that infiltrated into the lungs regardless of the type of virus used for infection. The one exception was a significant reduction in the number of B cells (CD45R^+^; Figure S2G) in TNF^-/-^ mice infected with ECTV^Rev.SECRET^ compared with those infected with ECTV^ΔCrmD^ or ECTV^WT^. TM mice were not included in the day 8 p.i. analysis of lung leukocytes. Results from the day 8 time-point suggested that the exacerbated lung pathology seen at this time cannot be attributed to quantitative changes in leukocyte numbers.

The flow cytometry experiments provided some key insights into the roles of TNF and CrmD pertaining to leukocyte recruitment to the lungs following ECTV infection. First, TNF signaling is critical for leukocyte recruitment into the lungs. Second, in WT mice, the SECRET domain plays a key role in leukocyte recruitment into the lungs in the first 4 days of infection. Third, in TM mice, interaction between mTNF and the TNF-binding domain of CrmD increased leukocyte numbers in the lungs of infected mice whereas the presence of the SECRET domain alone had no effect. As TM mice do not produce sTNF, TNFRI and TNFRII, the lack of any effect suggested that endogenous TNF signaling was necessary for SECRET to mediate its activity in relation to leukocyte recruitment into the lungs.

### vTNFRs dampen inflammation *in vitro* through reverse signaling

The binding of sTNFR to mTNF on a cell can induce reverse signaling *in vitro* and *in vivo* (Horiuchi et al., 2010; Juhasz et al., 2013; Watts et al., 1999) to dampen inflammation. We reasoned that CrmD and other OPV-encoded vTNFRs, which are known to bind mTNF (Pontejo et al., 2015b), could also potentially reverse signal through mTNF and downregulate the expression of inflammatory genes and as a consequence dampen inflammation in the lungs of infected hosts. Lipopolysaccharide (LPS) stimulation induced both sTNF and mTNF in the murine macrophage cell line RAW264.7 whereas it only induced mTNF in BMDM from TM mice (Figure S3). We used BMDM from TM mice stimulated with LPS to induce mTNF and inflammatory gene expression, and subsequently treated the cells with vTNFR We used the Mouse Signal Transduction Pathway Finder PCR array that screened for 84 genes (Figure S4; Table S3; (Benson et al., 2011)). If vTNFRs dampened inflammation, we expected to see the downregulation of mRNA transcripts for some LPS-induced inflammatory genes. We used recombinant CrmD from ECTV, CrmB, CrmC and CrmD from cowpox virus (CPXV) and CrmB from VARV expressed in the baculovirus system and purified as described (Pontejo et al., 2015a).

LPS stimulation upregulated the mRNA for 22 of the 84 genes that were screened (Figure S4). Treatment with Crm proteins downregulated 18 of the 22 mRNA upregulated by LPS (Table 1). Individual Crm proteins had varying effects on downregulating mRNA levels, with CPXV CrmD downregulating the greatest number of mRNA transcripts induced by LPS. All of the Crm proteins significantly reduced IL-1α, CCL2 and IL-2Rα mRNA levels following reverse signaling (Table 1). The pathways that were most affected by reverse signaling such as NF-κB and JAK-STAT cytokine signaling pathways are involved in inflammatory processes. Crm proteins also modulated those genes that were downregulated by LPS stimulation (Table S4) and these are represented by several signaling pathways including stress, NF-kB, CREB, Wnt and PIA3K/AKT pathways.

**Table 1.**
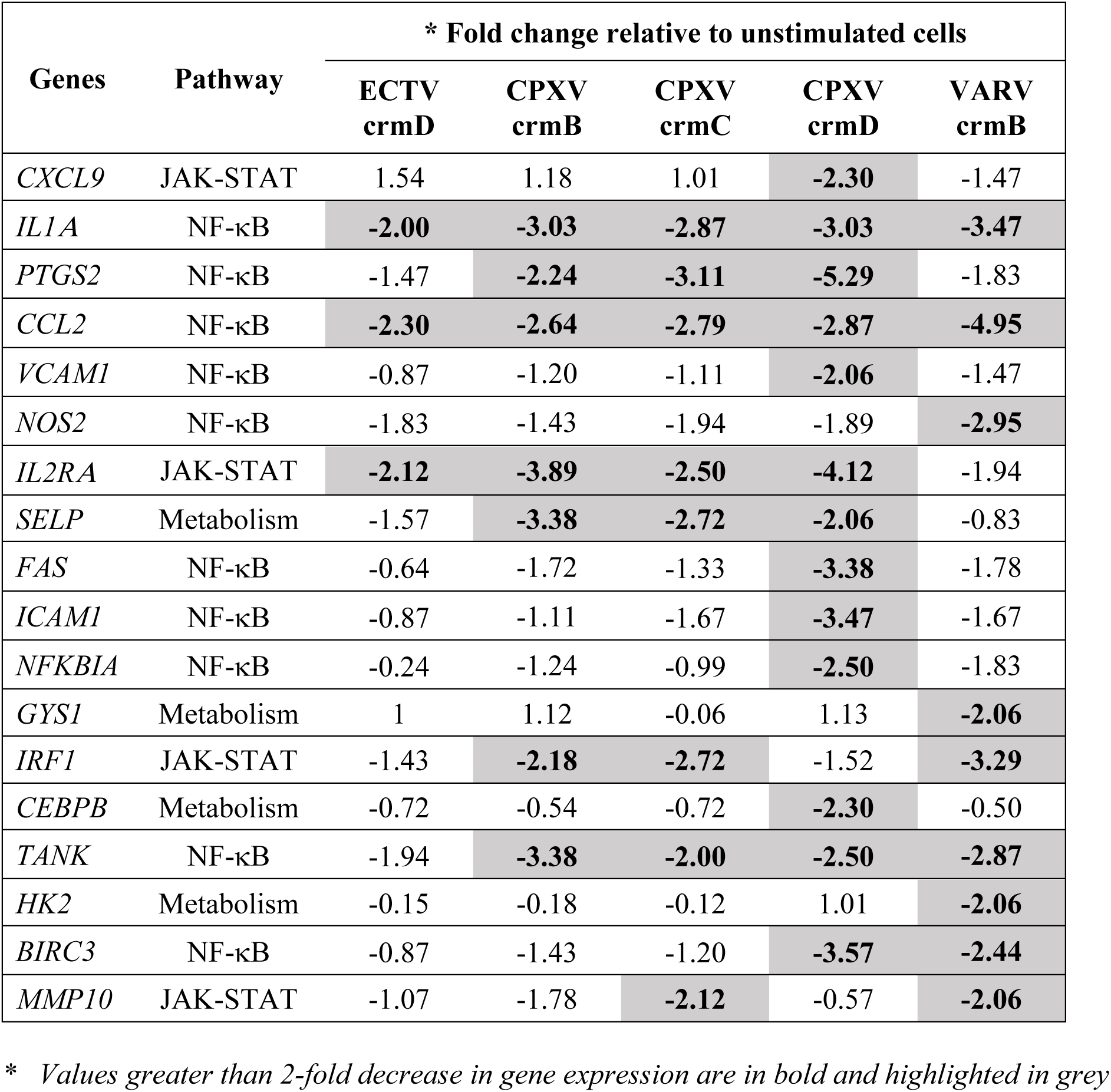
Genes upregulated by LPS stimulation and modulated following reverse signalling with vTNFRs

### Absence of CrmD results in dysregulated expression of pro-inflammatory mediators

Reverse signaling through mTNF by vTNFRs *in vitro* in LPS-stimulated BMDM cultures indicated that among others, the NF-κB and JAK-STAT signaling pathways were downregulated by vTNFRs. LPS stimulation of BMDM followed by vTNFR treatment may not be a true reflection of changes in mRNA expression that can occur in the various cells present in an organ as complex as the lung during viral infection, where many cell types produce TNF (Kels et al., 2019) and can thus be subjected to reverse signaling. We therefore determined changes in mRNA transcripts in lung tissue of WT, TNF^-/-^ and TM mice at day 9 p.i. using qRT-PCR. We also measured the protein levels of some of the cytokines in separate groups of infected WT mice.

The cytokine responses to ECTV^WT^ infection in WT and TM lungs were similar whereas the levels of most mRNA transcripts were significantly higher in TNF^-/-^ lungs compared to WT or TM lungs (Figures 5A-J). The absence of CrmD in ECTV^ΔCrmD^-infected WT mice led to significant increases in levels of mRNA transcripts for all mediators compared to those measured in lungs of WT mice infected with ECTV^WT^. In comparing mRNA transcript levels in lungs of WT mice infected with either mutant virus, three types of responses were apparent in the presence of the SECRET domain in ECTV^Rev.SECRET^-infected mice compared to those infected with ECTV^ΔCrmD^ infection. First, IL-1β, IL-12p40 and TGF-β mRNA levels were reduced compared to those induced by ECTV^ΔCrmD^ infection. Second, mRNA levels of IL-6 and IL-10 were increased significantly. Third, mRNA levels of TNF, IL-1α, IFN-γ, CCL2 and NOS2 were similar but were nonetheless significantly higher than levels induced by ECTV^WT^.

**Figure 5.**
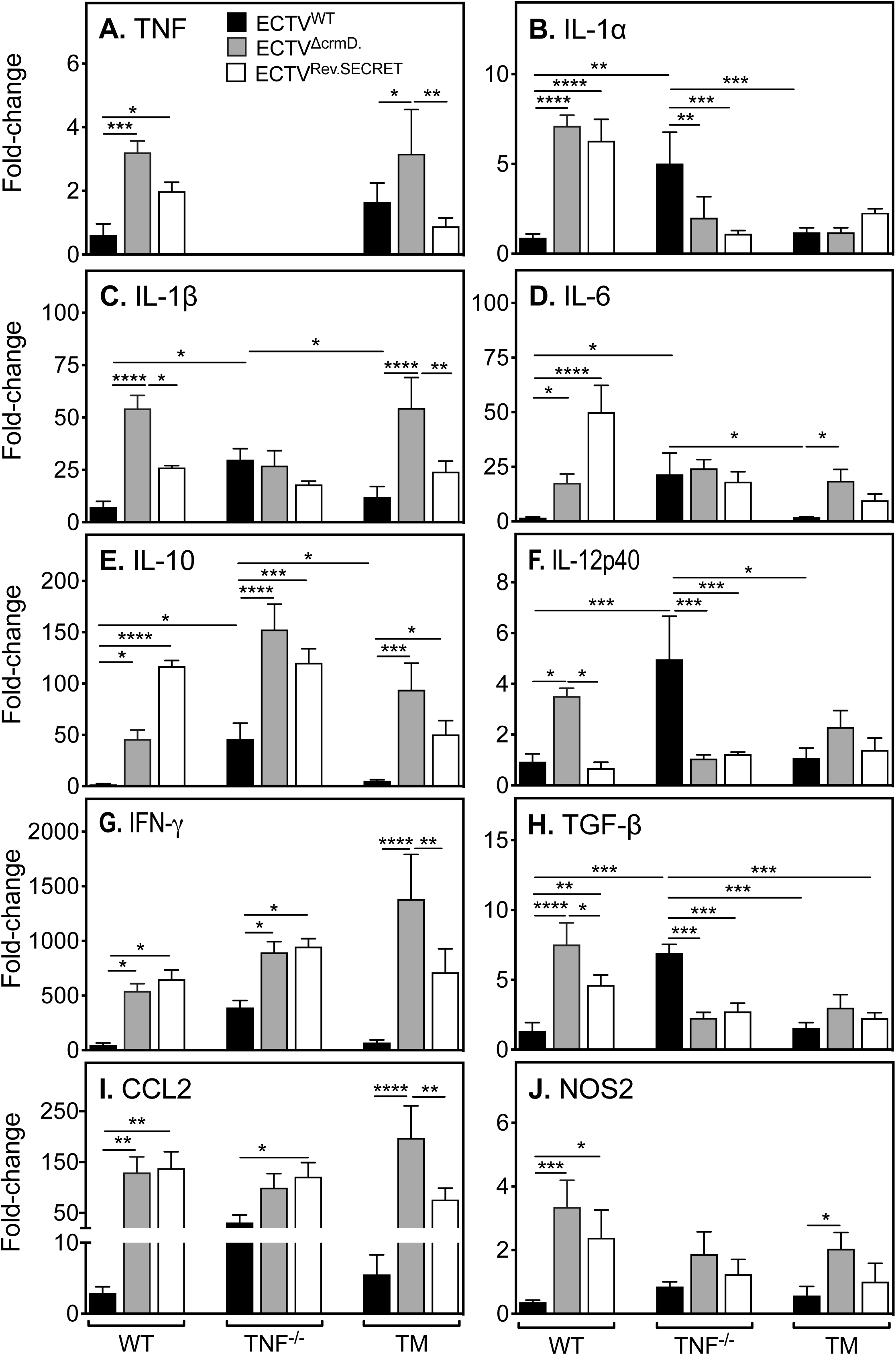
Deletion of CrmD significantly upregulates levels of mRNA transcripts for inflammatory mediators in virus-infected lungs. Groups of WT, TNF^-/-^ and TM mice (5 mice/group) were infected i.n. with 25 PFU ECTV^WT^, ECTV^ΔCrmD^ or ECTV^Rev.SECRET^. At day 9 p.i., mice were sacrificed and lungs were collected. RNA was isolated from lungs and qRT-PCR was used to determine levels of mRNA transcripts for (A) TNF, (B) IL-1α, (C) IL-1β, (D) IL-6, (E) IL-10, (F) IL-12p40, (G) IFN-γ, (H) TGF-β and (I) CCL2 and (J) NOS2. UBC was used as the reference gene. The expression levels of each gene in infected mice are relative to the expression levels in naive mice. Data are expressed as means ± SEM. Statistical analysis was performed using two-way ANOVA followed by Tukey’s multiple comparisons tests, where *, p < 0.05; **, p < 0.01; ***, p < 0.001. Data shown are from one of two separate experiments with comparable outcomes.

TNF^-/-^ mice did not express mRNA for TNF as expected but infection with ECTV^WT^ induced significantly higher levels of several inflammatory mediators compared to levels in WT or TM mice. The only exception was NOS2 mRNA, whose levels were similar to those in WT and TM mice. IL-1β, IL-6 and NOS2 mRNA levels in TNF^-/-^ mice infected with any one of the 3 viruses were comparable. Some mRNA transcripts in TNF^-/-^ mice infected with either of the mutant viruses changed significantly. IL-10, IFN-γ and CCL2 levels increased whereas IL-1α, IL-12p40 and TGF-β levels decreased regardless of whether both domains were absent or only the SECRET domain was present. This finding suggested that the changes in mRNA expression in the absence of TNF might have been due to interactions of the TNF-binding domain of CrmD with LT-α and/or LT-β.

In TM mice, CrmD deficiency led to increases in levels of expression of a number of mediators, except IL-1α, IL-12p40 and TGF-β. TNF and IL-1β levels were significantly higher in the ECTV^ΔCrmD^-infected group compared to ECTV^WT^- or ECTV^Rev.SECRET^-infected mice. Similarly, IL-6 and IL-10 levels were higher in the absence of CrmD, reduced when only the SECRET domain was present but were clearly higher than in ECTV^WT^-infected animals. NOS2 mRNA transcripts were only increased in the lungs of mice infected with ECTV^ΔCrmD^.

Both the TNF-binding and SECRET domains differentially regulated mRNA transcripts for a number of inflammatory genes in the 3 different strains of mice. In WT animals, the TNF-binding domain was particularly important for dampening TNF, IL-1α, IL-1β, TGF-β, CCL2, IFN-γ and NOS2 whereas the SECRET domain appeared critical for induction of IL-6 and IL-10. In TM mice, the presence of the SECRET domain alone reduced levels of CCL-2, IFN-γ, IL-1β, IL-6, IL-10 and TNF compared to levels in the absence of both the TNF-binding and SECRET domains.

We measured the levels of some cytokines by ELISA in lung homogenates of WT mice infected with ECTV^WT^, ECTV^ΔCrmD^ or ECTV^Rev.SECRET^ (Figure S5). The absence of CrmD in ECTV^ΔCrmD^- infected mice increased levels of TNF, IL-10 and CCL2 whereas the presence of only the SECRET domain in ECTV^Rev.SECRET^-infected mice significantly increased levels of IL-6 and IL-10, consistent with the gene expression data (Figure 5). CrmD deficiency induced the highest levels of IL-1β and IL-12p40 mRNA transcripts however the SECRET domain alone was sufficient to induce significantly higher levels of these proteins. Furthermore, while the absence of CrmD significantly increased levels of mRNA transcripts for IFN-γ, the protein levels were nonetheless comparable in lung homogenates from mice infected with ECTV^WT^, ECTV^ΔCrmD^ or ECTV^Rev.SECRET^. These differences might have been due to post-transcriptional regulation of cytokine gene expression, and/or the cytokine milieu in the lungs of the different strains of mice infected with the different viruses may have impacted on cytokine protein translation.

### Blockade of IFN-γ, IL-6, IL-10R or TNF reduces weight loss and disease and increases survival rates

The preceding data established that infection of mice with mutant ECTV lacking both the TNF-binding and SECRET domains or lacking only the TNF-binding domain resulted in increased production of a number of inflammatory cytokines, possibly contributing to an increase in morbidity and mortality. We postulated that the *in vivo* blockade of some of those cytokines might reduce the severity of the clinical symptoms and pathology and result in the recovery of mice infected with ECTV^ΔCrmD^ or ECTV^Rev.SECRET^.

WT mice infected with ECTV^ΔCrmD^ or ECTV^Rev.SECRET^ were treated with mAb against IFN-γ, IL-6, IL-10R or TNF, and treatment continued every alternate day until day 20 p.i. and surviving animals were sacrificed on day 21. We selected day 7 p.i. to start cytokine blockade therapy as this is the time when animals start losing weight, have relatively higher clinical scores and lung pathology starts to develop. Control groups were treated with an isotype-matched rat IgG mAb. All animals were monitored daily for clinical signs of disease, weight loss and survival.

In ECTV^ΔCrmD^-infected WT mice, blockade of IFN-γ, IL-6, TNF or IL-10R with specific mAb significantly increased survival rates compared to the rat IgG isotype control mAb-treated group (Figure 6A). The control mAb-treated group exhibited 100% mortality by day 8 p.i. whereas 60% of animals treated with mAb against TNF or IL-10R were alive at day 21. Anti-IL-6 mAb treatment resulted in the survival of 80% of animals in the group whereas IFN-γ blockade rescued 100% of the mice. Individually, IFN-γ, IL-6, TNF and IL-10 were necessary and sufficient to cause significant morbidity and mortality due to lung pathology but the blockade of just one of the cytokines was sufficient to protect 60-100% of the mice.

**Figure 6.**
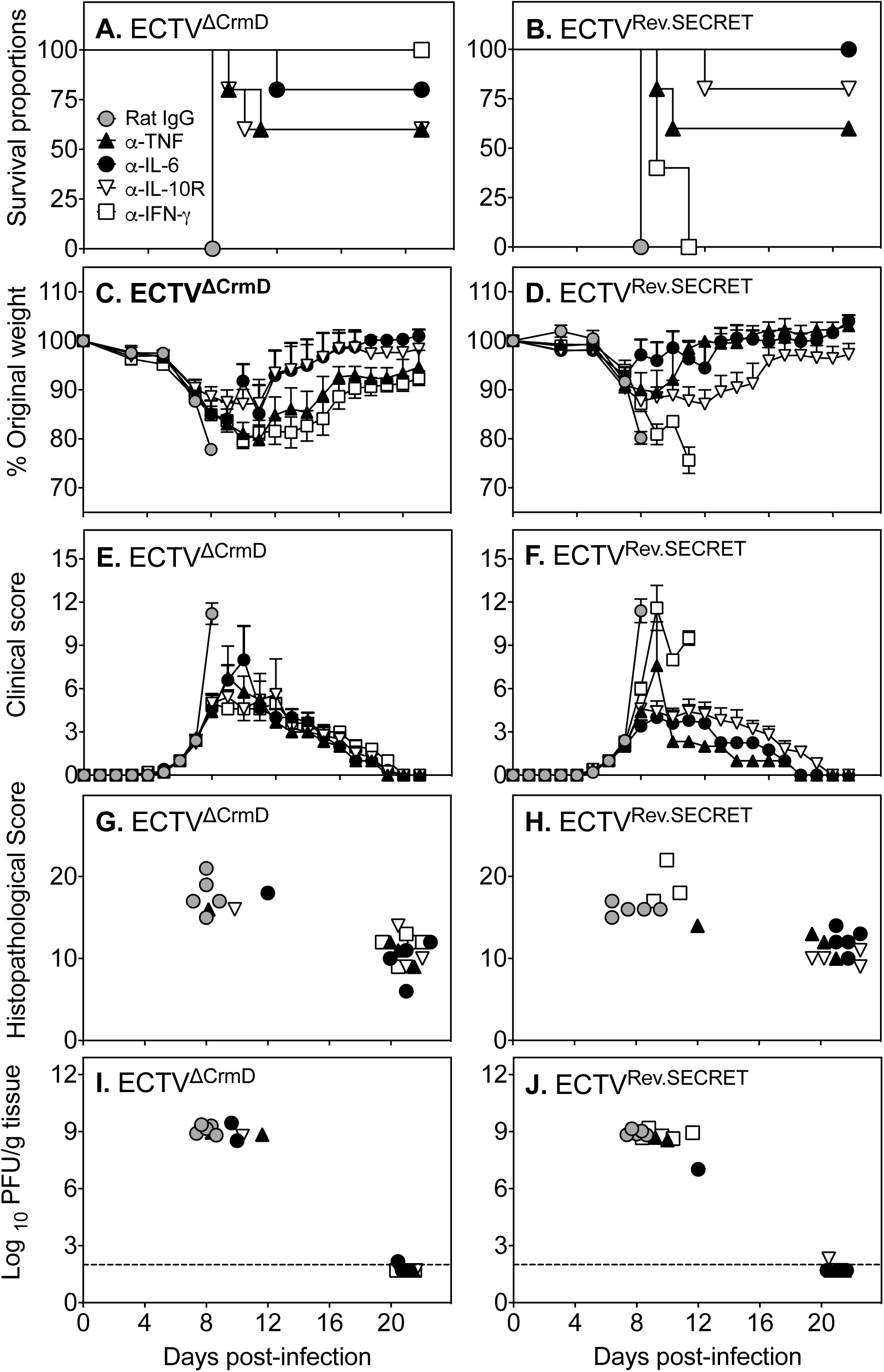
Blockade of TNF, IFN-γ, IL-6 or IL-10R protects against mutant virus infection. Groups of WT female mice (5 mice/group) were infected i.n. with 25 PFU of ECTV^ΔCrmD^ or ECTV^Rev.SECRET^. On days 7, 9, 11, 13, 15, 17, 19 and 21 p.i., individual groups were treated with isotype control rat IgG mAb (control mAb) or specific mAb against TNF (α-TNF), IL-6 (α-IL-6), IL-10R (α-IL-10R) or IFN-γ (α-IFN-γ) at 500 µg i.p. Mice were monitored daily and euthanized if they had lost ≥25% of their original weight or were severely moribund at any time during the course of infection. All surviving animals were euthanized on day 22 p.i. (A, B) Survival, (C, D) weight loss, (E, F) clinical scores, (G, H) histopathological scores, and (I, J) viral load in mice infected with ECTV^ΔCrmD^ or ECTV^Rev.SECRET^. For A and B, statistical analysis was done using the Log-rank (Mantel-Cox) test, p < 0.01 in survival proportions of mice infected with ECTV^ΔCrmD^ or ECTV^Rev.SECRET^ and treated with anti-cytokine mAb compared to mice treated with control mAb. For C, D, E and F, significance was obtained using two-way ANOVA followed by Dunnett’s post-tests compared to the means of control mAb-treated groups at day 8 p.i. For C and D, in the ECTV^ΔCrmD^-infected groups, p < 0.001 when control mAb treated group is compared with α-IL-6- or α-IFN-γ-treated groups and p < 0.0001 when control mAb treated group is compared with α-TNF or α-IL-10R treated groups. In the ECTV^Rev.SECRET^-infected groups, p < 0.0001 when control mAb treated group is compared with α-IL-6- or α-IFN-γ-treated groups and p < 0.01 when control mAb-treated group is compared with α-TNF or α-IL-10R treated groups. For E and F, in the ECTV^ΔCrmD^- and ECTV^Rev.SECRET^-infected groups, p < 0.0001 when control mAb-treated group is compared with α-IL-6-, α-IFN-γ-, α-TNF- or α-IL-10R-treated groups. For G, H, I and J, data sets for individual mice are presented as the results come from different days for each of the groups. For that reason, statistical analysis was not done. Data shown are from one experiment.

ECTV^Rev.SECRET^-infected mice also succumbed to infection by day 8 p.i. (Figure 6B). Survival rates significantly increased from 0% to 60% and from 0% to 80% after treatment with anti-TNF or anti-IL-10R, respectively. There was 100% survival in the group treated with anti-IL-6 (Figure 6B). Thus, blockade of IL-6, TNF or IL-10R resulted in similar outcomes in animals infected with ECTV^ΔCrmD^ or ECTV^Rev.SECRET^. However, the only exception was anti-IFN-γ treatment of ECTV^Rev.SECRET^-infected mice, which had minimal effects on the outcome and all animals succumbed to infection. This finding is in complete contrast to ECTV^ΔCrmD^-infected mice which were fully protected from anti-IFN-γ treatment. Thus, IFN-γ, IL-6, TNF and IL-10 are responsible for the pathology in the absence of the TNF-binding and SECRET domains in ECTV^ΔCrmD^- infected mice whereas in the presence of only the SECRET domain in ECTV^Rev.SECRET^-infected mice, IFN-γ is neither necessary nor sufficient to cause pathology.

Weight loss (Figures 6C and 6D) and clinical scores (Figures 6E and 6F) were also markedly lower after anti-cytokine treatment compared to the control mAb-treated groups. Mice treated with anti-cytokine mAb regained weight beginning at day 12 and those mice that recovered had similar clinical scores from this time. Improvements in weight loss and clinical scores correlated with the survival rates, as evidenced by body weight recovery and the absence of any clinical signs and symptoms in mice that were alive at day 21 p.i.

The histopathological scores (Figures 6G and 6H) generated from lung histology sections (Figure 7) were highest in those animals that had the highest clinical scores, lost significant body weights and succumbed to mousepox compared to those animals that had survived. The viral load was very high in those animals that died early between days 8 and 12, but was at or below the limit of detection in those animals that recovered and which were euthanized at day 21 (Figures 6I and 6J). Of particular interest is the outcome of IFN-γ blockade in mice infected with ECTV^ΔCrmD^ or ECTV^Rev.SECRET^. IFN-γ has a well-established antiviral role in the mousepox model (Karupiah et al., 1993; Panchanathan et al., 2005). In mice infected with ECTV^ΔCrmD^ and treated with anti-IFN-γ, viral load was below the limit of detection and the histopathological scores were substantially reduced. In contrast, in mice infected with ECTV^Rev.SECRET^ and treated with anti-IFN-γ, the viral load and histopathological scores were comparable with those of control mAb-treated mice. Thus, excessive IFN-γ production in mice infected with ECTV^ΔCrmD^, when both TNF-binding and SECRET domains are absent, results in significant lung pathology. However, excessive IFN-γ production in mice infected with ECTV^Rev.SECRET^, when only SECRET domain is present, is not sufficient to cause pathology.

**Figure 7.**
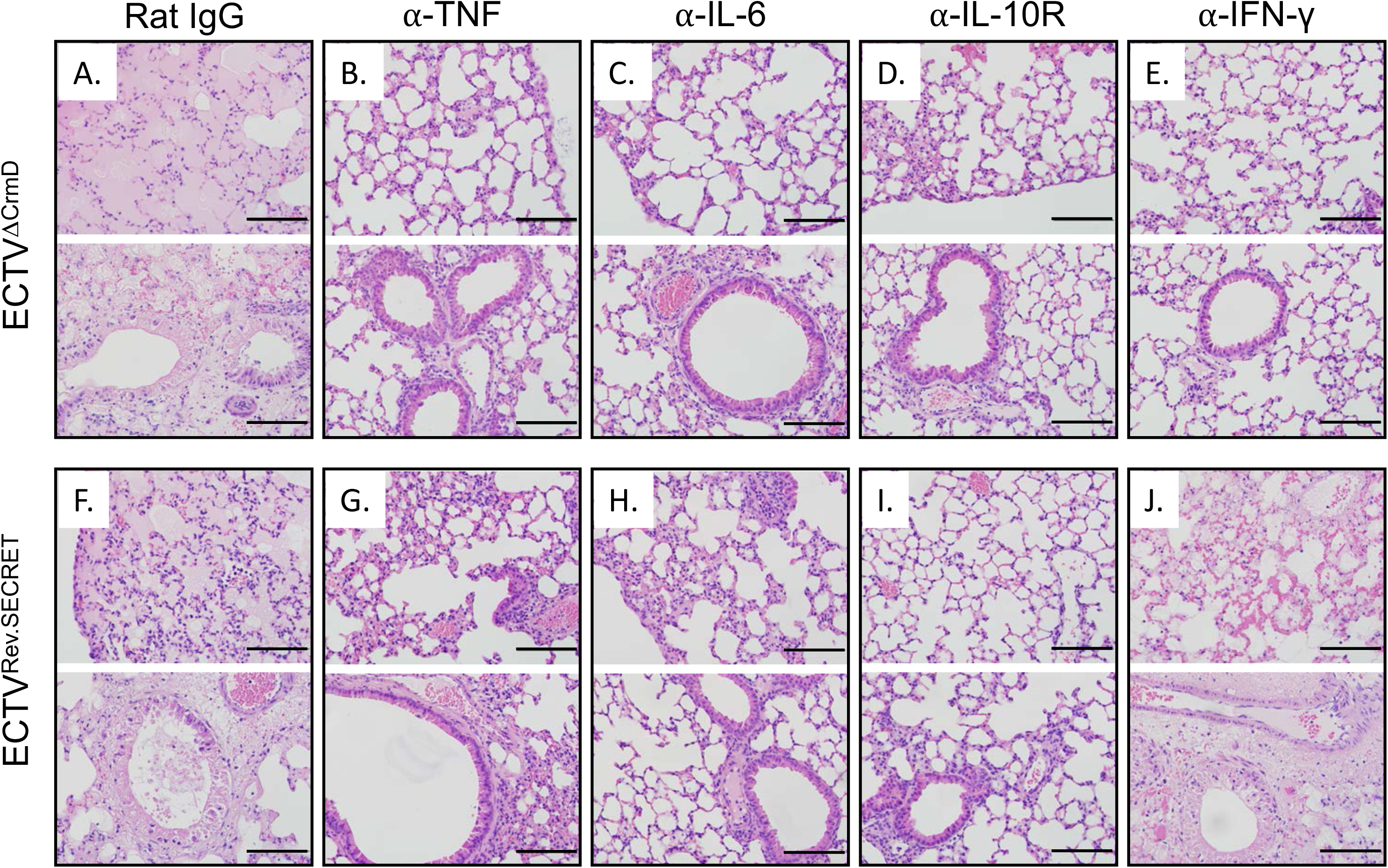
Reduction in inflammation and lung pathology in ECTV^ΔCrmD^- or ECTV^Rev.SECRET^- infected mice treated with mAb to TNF, IFN-γ, IL-6 or IL-10R. H&E-stained histological sections from the experiment described in Figure 6. Mice were ethically euthanized if they lost ≥25% of their original weight or were severely moribund based on the clinical scores. Lungs were fixed, paraffin-embedded, sectioned and stained with H&E. Slides were examined on all fields at 400x magnification. Representative sections are from ECTV^ΔCrmD^- or ECTV^Rev.SECRET^-infected mice treated with control mAb (A, F), α-TNF (B, G), α-IL-6 (C, H), α-IL-10R (D, I), or α-IFN-γ (E, J). Bars correspond to 100 µm.

The results indicate that the severity of ECTV infection seen in the absence of both the TNF-binding and SECRET domains or only the TNF-binding domain was due to the excessive production of a number of pro-inflammatory cytokines which contributed to lung pathology, morbidity and mortality. Blockade of the functions of specific cytokines reduced morbidity and lung pathology and improved survival rates, some up to 100%.

## Discussion

OPV encode homologs of mammalian cytokine and chemokine receptors that can dampen, evade or subvert the host immune response and influence the outcome of infection (Alcami, 2003; Alcami and Koszinowski, 2000; Seet et al., 2003; Stanford et al., 2007). Such a strategy provides an advantage to the virus, allowing successful replication, transmission to other hosts and even persistence within the host (Sakala et al., 2015b). As viruses have co-evolved with their hosts and virus-encoded molecules can be host species specific, we utilized ECTV, a natural mouse pathogen, to address the function of CrmD in C57BL/6 mice that are normally resistant to infection with this virus. Both VARV and ECTV are very closely related OPV that cause respiratory infection and as mousepox is a robust surrogate model for smallpox. The findings with mousepox will be relevant to smallpox and specifically to VARV-encoded CrmB, a vTNFR with TNF/LT binding properties for human TNF/LT similar to those of CrmD for mouse TNF/LT, and also containing a SECRET domain with similar binding properties for chemokines (Alejo et al., 2006; Pontejo et al., 2015a). Respiratory infection of C57BL/6 mice with ECTV^WT^ results in morbidity but animals completely recover whereas infection with ECTV^ΔCrmD^ or ECTV^Rev.SECRET^ caused significant morbidity and 100% mortality. In the ECTV-resistant C57BL/6 mice, CrmD actually provided an advantage to the host. The TNF-binding domain of CrmD not only reduces the bioavailability of host TNF but it also dampens inflammatory cytokine production by “reverse signaling” through mTNF. The combined results from both *in vivo* experiments in ECTV^WT^-, ECTV^ΔCrmD^- or ECTV^Rev.SECRET^ -infected mice and *in vitro* results from BMDM treated with Crm proteins from various OPV together revealed that the binding of CrmD to mTNF significantly reduces the levels of TNF production, in addition to that of a number of other cytokines that can mediate inflammatory or regulatory functions.

Previous studies on ECTV-encoded cytokine binding proteins (bp) such as IFN-γ-bp; (Sakala et al., 2015a), IFN-α/β-bp (Sakala et al., 2015a; Xu et al., 2008) and IL-18-bp (Sakala et al., 2015a) revealed that infection of ECTV-susceptible BALB/c mice with deletion mutant viruses was attenuated, some more than others. These mutant viruses were also attenuated in C57BL/6 mice but the effects were far more dramatic in BALB/c mice. Indeed, some of us recently reported that infection of BALB/c mice with ECTV^ΔCrmD^ attenuated the infection to the extent that animals fully recovered from an otherwise lethal infection (Alejo et al., 2018). When infected with ECTV^WT^, BALB/c mice respond with a weak inflammatory response and produce very little TNF or other inflammatory cytokines (Chaudhri et al., 2004). However, when BALB/c mice were infected with ECTV^ΔCrmD^, the inflammatory and cell-mediated immune responses were augmented resulting in effective virus control and survival (Alejo et al., 2018). Both the SECRET and TNF-binding domains were found to be required for CrmD to mediate its function fully. The TNF-binding domain of CrmD neutralized TNF, whereas the SECRET domain reduced leukocyte recruitment, resulting in a reduction in the capacity of the host to generate an effective antiviral immune response (Alejo et al., 2018).

In the C57BL/6 mouse model of ECTV infection used in the present study, expression of a soluble cytokine inhibitor protected the host from virus-induced immunopathology, instead of increasing virulence. We found that deletion of CrmD from ECTV significantly increased morbidity and mortality in infected C57BL/6 mice, which are normally resistant to ECTV^WT^ infection. It is interesting to note that deletion of the secreted IL-1β receptor encoded by vaccinia virus also caused significant detrimental effects after i.n. infection of BALB/c mice, in this case with increased fever and accelerated weight loss and mortality (Alcami and Smith, 1992, 1996). We have found that CrmD is critical for dampening lung inflammation in ECTV-resistant C57BL/6 mice and that its function requires host TNF. However, in contrast to BALB/c mice, C57BL/6 mice infected with ECTV^ΔCrmD^ succumbed to mousepox due to significant lung inflammation and pathology. The increased susceptibility was not due to increased lung viral load as titers of WT and mutant viruses were comparable across the WT, TNF^-/-^ and TM mice. C57BL/6 WT mice produce significantly higher levels of TNF compared to BALB/c mice (Chaudhri et al., 2004). The absence of CrmD in WT and TM mice infected with ECTV^ΔCrmD^ resulted in significantly increased levels of TNF and a number of other inflammatory cytokines including IL-6, IL-10, TGF-β and IFN-γ. The dysregulated cytokine responses were contemporaneous with weight loss, higher clinical scores, reduced leukocyte recruitment to the lungs, exacerbated lung pathology and 100% mortality in both WT and TM mice. WT mice in particular demonstrated clear increases in inflammatory cytokine responses but significantly reduced leukocyte recruitment into the lungs when both the TNF-binding and SECRET domains were absent. The presence or absence of the SECRET domain when the TNF-binding domain was absent did not affect clinical scores, weight loss or survival of WT and TM mice. However, cellular infiltrates were increased in ECTV^Rev.SECRET^-infected lungs compared to ECTV^ΔCrmD^-infected lungs. It is apparent that the TNF-binding domain of CrmD reduced leukocyte infiltration into the lungs whereas the SECRET domain contributed to leukocyte recruitment. The findings in C57BL/6 mice are in sharp contrast to those in BALB/c mice (Alejo et al., 2018). As the SECRET domain has chemokine-binding activity and can bind to the chemokines CCL-25, CCL-27, CCL-28, CXCL-11, CXCL-12β, CXCL-13 and CXCL-14 (Alejo et al., 2018), we had predicted that the SECRET domain would act in a similar manner in the C57BL/6 strain. It is likely that absence of the TNF-binding domain of CrmD in ECTV^Rev.SECRET^-infected mice, which resulted in excessive production of a number of inflammatory cytokines, overcame the inhibitory effects of the SECRET domain on leukocyte recruitment to the lungs. It is also likely that virus inoculation through different routes, i.e. the i.n. route in C57BL/6 mice as in the current study and through the subcutaneous route in the foot-pad in BALB/c mice (Alejo et al., 2018) may induce a distinct pattern of chemokine expression and influence leukocyte recruitment.

The C57BL/6 TNF^-/-^ strain is highly susceptible to respiratory ECTV^WT^ infection and dies from serious lung pathology (Kels et al., 2019). Apart from binding to TNF, the TNF-binding domain of CrmD can also interact with LT-α and LT-β and use of TNF^-/-^ mice helped to dissociate the effects of CrmD binding to TNF from those of binding to LT-α or -β. Unlike the significant effects seen in WT mice, infection of TNF^-/-^ mice with ECTV^ΔCrmD^ or ECTV^Rev.SECRET^ did not make any difference to weight loss, clinical scores, leukocyte recruitment, viral load, lung pathology or survival. These findings suggest that the major biological functions of the TNF-binding domain of CrmD are mediated through binding to TNF. Nonetheless, there were clearly some mRNA transcripts for cytokines like IL-1α, IL-12p40 and TGF-β that were down-regulated and others like IFN-γ and CCL-2 that were upregulated in TNF^-/-^ mice. Those changes in gene expression most likely reflect the potential interactions of the TNF-binding domain with LT-α and/or LT-β but they did not impact on the outcome of infection.

Although the presence or absence of the SECRET domain when the TNF-binding domain was absent did not affect clinical scores, weight loss or survival of WT and TM mice, there were clear differences in the requirement for IFN-γ in causing pathology or controlling virus load. IFN-γ has a well-established antiviral role in the mousepox model (Karupiah et al., 1993; Panchanathan et al., 2005) but excessive production can also cause lung pathology (Kels et al., 2019). In mice infected with ECTV^ΔCrmD^ and treated with anti-IFN-γ, viral load was below the limit of detection and the histopathological scores were substantially reduced. In contrast, in mice infected with ECTV^Rev.SECRET^, viral titers and histopathological scores were comparable with those of control mAb-treated mice. Thus, excessive IFN-γ production in mice infected with ECTV^ΔCrmD^, i.e. when both the TNF-binding and SECRET domains were absent, resulted in significant lung pathology and possibly ineffective virus control. However, excessive IFN-γ production in mice infected with ECTV^Rev.SECRET^, which expressed only the SECRET domain, was not sufficient to cause pathology. While the precise reasons for these differences warrant further investigation, the local cytokine and chemokine milieu, and the number, nature and activation status of infiltrating leukocytes under conditions of infection with the 2 different viruses may have played a significant role. For instance, cellular infiltrates were increased in ECTV^Rev.SECRET^-infected lungs compared to ECTV^ΔCrmD^-infected lungs. Furthermore, at least in WT mice, presence of the SECRET domain alone induced high levels of IL-6 and IL-10. Excessive IL-6 or IL-10 production, as a consequence of TNF deficiency or excessive TNF production, during respiratory ECTV infection causes significant lung pathology (Kels et al., 2019). It is evident that both the TNF-binding and SECRET domains need to work cooperatively to dampen pathology in C57BL/6 mice in order to provide an advantage to the host. In sharp contrast, in BALB/c mice, both the domains are important to effectively dampen inflammation and the immune response in order to give an advantage to the virus.

It is interesting to note that under conditions of excessive TNF production, as is the case in C57BL/6 mice infected with the CrmD mutant ECTV, and TNF deficiency, as is the case with infection of TNF^-/-^ mice with ECTV (Kels et al., 2019), an overlapping set of cytokines is dysregulated. It is well established that excessive TNF production involving NF-κB activation can induce the production of other inflammatory cytokines, including IL-6 and IL-10, which in turn can activate signal transducer and activator of transcription (STAT) 3 (Zhong et al., 1994). Given that NF-κB and STAT3 signaling pathways are closely intertwined and regulate an overlapping group of target genes including those associated with inflammation (Goldstein et al., 2017; Kasembeli et al., 2018; Ma et al., 2017; Oeckinghaus et al., 2011), perturbations in the NF-κB signaling pathway will disrupt or dysregulate the STAT3 signaling pathway as well.

In conclusion, the outcome of infection of C57BL/6 versus BALB/c mice with vTNFR mutant viruses would be determined not only by a balance between the host’s ability to produce TNF and the virus’ ability to neutralize its effects but also how CrmD signaling through mTNF affects production of other inflammatory cytokines. Based on the mousepox model, we would speculate that in humans, VARV-encoded CrmB may also protect against lung pathology in those individuals capable of generating potent inflammatory responses but otherwise contribute to increased susceptibility in those individuals not capable of generating an effective immune response.

## Supporting information

Suppplementary Figures

## Acknowledgements

This work was supported by a grant from the National Health and Medical Research Council of Australia to G.K. and G.C. (grant ID APP 1007980). The laboratory of A. Alcami was funded by the Spanish Ministry of Economy and Competitiveness and European Union (European Regional Development’s Funds, FEDER) (Grant SAF2015-67485-R). The funders had no role in study design, data collection and interpretation, or the decision to submit the work for publication. We would like to thank Professor Jane Dahlstrom of the Australian National University Medical School and the Canberra Hospital, Australian Capital Territory, Australia for helping us to develop the scoring system for the histopathology of lung sections. We also thank Dr. Jonathon D. Sedgwick, Boehringer Ingelheim Pharmaceuticals Inc., Ridgefield, USA for the gift of the TNF^-/-^ and mTNF^Δ/Δ^ mice. We acknowledge the assistance of staff at the Australian National University Phenomics Facility for animal breeding and the John Curtin School of Medical Research Microscopy and Flow Cytometry Research Facility.

## Author contributions

Conceptualization, Z.A.R., M.J.T.K., E.N., A. Alcami, G.C. and G.K.; Methodology, Z.A.R., M.J.T.K., E.N., G.C. and G.K.; Investigation, Z.A.R., M.J.T.K., E.N.; Writing – Original Draft, G.C. and G.K.; Writing – Review & Editing, Z.A.R., M.J.T.K., E.N., P.P., A. Alejo, S.M.P. and A. Alcami, G.C. and G.K.; Funding Acquisition, A. Alcami, G.C. and G.K.; Resources, A. Alejo, S.M.P. and A. Alcami, G.C. and G.K.; Supervision, G.C. and G.K.

## Methods

### Mice

Six- to twelve-week old female WT, Tnf^tm1Jods^ (TNF^-/-^) (Korner et al., 1997), Tnf^tm2Jods^ (mTNF^Δ/Δ^) (Ruuls et al., 2001) and Tnf^tm2Jods^Tnfrsf1a^tm1Imx^Tnfrsf1b^tm1Imx^ (designated triple mutant, TM) mice on a C57BL/6 background were bred under specific pathogen-free conditions at the Australian Phenomics Facility, Australian National University, Canberra, Australia. WT mice express sTNF, mTNF TNFRI and TNFRII. TNF^-/-^ mice are deficient in sTNF and mTNF but express TNFRI and TNFRII. The mTNF^Δ/Δ^ mice express non-cleavable mTNF, TNFRI and TNFRII but not sTNF. Triple mutant (TM) mice express non-cleavable mTNF but not sTNF, TNFRI or TNFRII. We generated the TM mice by crossing the mTNF^Δ/Δ^ with TNFRI- and TNFRII-deficient mice (Peschon et al., 1998). All animal experiments were performed in accordance with protocols approved by the Animal Ethics and Experimentation Committee of the Australian National University (Protocol numbers A2011/011 and A2014/018) and the Animal Ethics Committee of the University of Tasmania (Protocol number A0016372).

### Cell lines

BS-C-1 (ATCC^®^ CCL-26^™^) cells were cultured in Eagle’s minimum essential medium (EMEM) supplemented with 2 mM L-glutamine (Sigma Aldrich), antibiotics (penicillin, 120 µg/mL, streptomycin, 200 µg/mL and neomycin sulphate), 1 mM 4-(2-hydroxyethyl)-1-piperazineethanesulfonic acid (HEPES) and 10% heat-inactivated fetal calf serum (FCS). This medium referred to as complete EMEM10. RAW264.7 (RAW 264.7 (ATCC^®^ TIB-71^™^) and NCTC clone 929 (L-929 cells; ATCC^®^ CCL-1^™^) cells were cultured in complete Dulbecco’s modified Eagle’s medium (DMEM10) with additional supplements of 1 mM sodium pyruvate, 1 mM non-essential amino acids.

### Viruses

The Naval strain of WT ECTV (ECTV^WT^), the CrmD deletion mutant (ECTV^ΔCrmD^) and the CrmD deletion mutant revertant expressing only the SECRET domain (ECTV^ΔCrmD.Rev.SECRET^ and abbreviated as ECTV^Rev.SECRET^ for simplicity) (Alejo et al., 2018) were used. The construction of the mutant and revertant viruses has been described in detail by Alejo et. al (2018). In data shown in Figure S1, the Moscow strain of ECTV (ATCC No. VR-1374) was used. Viruses were propagated in BS-C-1 cells and single-use aliquots were stored at −80°C as described elsewhere in detail (Chaudhri et al., 2018).

### Viral plaque assay for quantification of ECTV

To enumerate the number of virus plaque forming units (PFU) present in samples, a viral plaque assay was employed as previously described (Chaudhri et al., 2018; Scalzo et al., 2000). BSC-1 cells were seeded at 1 x 10^5^ cells per well in a sterile 12-well cell culture plate (Nunc, NY, USA) a day before the assay. Organs were aseptically removed from euthanized mice on selected days and immediately snap frozen on dry ice. Samples were then stored at −80°C until required. On the day of the assay, organs were thawed, weighed and 1mL of phosphate buffered saline (PBS) supplemented with 1 mM HEPES and 1% FCS was added to each sample. Organs were homogenized either with a polytron homogenizer (ProScientific Inc., Monroe, CT, USA) or the TissueLyser II (Qiagen No. 85300) at 25 Hz for 2 min twice. After disruption of the tissue, the mixture was sonicated at 100W power in three 8-second bursts in a cup sonicator to further break up clumps of virus (Branson Sonic Power Company, Danbury, CT, USA). To pellet unwanted organ debris, samples were centrifuged at 3000 x g for 5 min at 4°C. Six 10-fold serial dilutions of the supernatant were made and 100 µL added to confluent BSC-1 monolayers. The plates were then incubated at 37°C for 1 hour before 1 mL of overlay (1% methylcellulose (viscosity 400 centipoise, Spectrum Chemical Manufacturing corporation, NJ, USA), 2.5% FCS, 1 mM HEPES, 2 mM L-glutamine in EMEM) was added. Following 4 days of incubation at 34°C, the BSC-1 monolayers were stained with 200 µl 0.1% crystal violet to visualize the plaques. The viral titer was then calculated from the following equation:

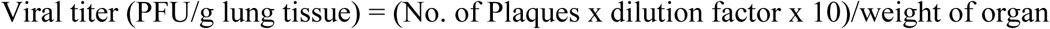

### Preparation of L929-cell conditioned medium (LCCM)

L-929 cells were grown in DMEM10 and maintained at 37°C in a 5% CO2 humidified incubator until cells reached confluency and the medium exhausted. The supernatant was then collected, passed through a 0.45 μM filter, and stored in 5-mL aliquots at −20°C. LCCM, a source of macrophage-colony stimulating factor (M-CSF), was used for generation of BMDM.

### Generation of BMDM

To determine whether the TNF-binding or SECRET domains are essential for ECTV replication *in vitro*, BMDM from WT, TNF^-/-^ and TM mice were isolated and cultured as follows. Mice were euthanized and their femurs and tibia were collected aseptically. Excess muscles were removed and clean bones were placed into cold PBS. The ends of each bone were removed and a 25-G needle attached to a 10 mL syringe filled with cold PBS was inserted into the bone marrow cavity and flushed with 2-5 mL fluid until it appeared white. Medium containing the BM cells was collected in 50 mL tubes, clumps dispersed using an 18-G needle attached to a 10 mL syringe and cell suspensions were filtered through a 70 μm nylon cell strainer. The cells were washed twice (centrifugation for 5 min at 400 x g at 4°C) and resuspended in complete DMEM10. Total BM progenitor cells were counted and adjusted to 2 x 10^6^ cells/mL of complete DMEM10 containing 10% L-929 cell conditioned medium (LCCM). For each 25 cm2 flask, 2 x 10^7^ cells in 10 mL were seeded and incubated at 37°C and 5% CO_2_ humidified incubator. After incubation for 6-18 h, non-adherent cells were transferred to a 75 cm2 flask and an additional 20 mL medium was added to each flask. These cells were cultured for 4 days allowing them to adhere and multiply; the growth medium was changed every 2 days. After a further 2-3 days of incubation, the cells were ready for harvesting. Mature macrophages were gently scraped in a unidirectional manner with a sterile cell scraper into the medium, the cells centrifuged at 200 x g for 5 min at 4°C and resuspended in 10% complete DMEM.

### Virus replication kinetics in BMDM

Approximately 2 x 10^6^ cells in 1 mL DMEM10 were seeded into wells of 12-well cell culture plate (Nunc, NY, USA). Two hours later after the cells had adhered, the medium was removed and replicate wells were infected with the CrmD mutant or WT parent virus at a multiplicity of infection (MOI) of 1 in a 100 μL volume of DMEM containing 1% FCS and allowed to adsorb for 1 h. Next, 1 mL PBS was added to each well gently and aspirated to remove non-adsorbed virus followed by the addition of 1 mL complete DMEM10 to wells and plates were incubated at 37°C in a 5% CO_2_ humidified incubator. Cells and media in each well were harvested at 0 (1 h post-adsorption), 6, 12, 24 and 48 h p.i. and stored at −80°C. All frozen samples were processed at the same time to measure virus titers using viral plaque assay as described in detail elsewhere (Chaudhri et al., 2018).

### Determination of total protein concentration in homogenized lungs

Lungs were collected and homogenized in 1 mL RIPA buffer (150 mM NaCl, 0.1% Triton X-100, 0.5% sodium deoxycholate, 50 mM Tris-HCl pH 8.0) with 0.1 μL Protease Inhibitor Cocktail VI (Astral Scientific No. P-1515) using the TissueLyser II (Qiagen No. 85300) at 25 Hz for 2 min twice. Lung homogenates were incubated on ice at 4°C and rocked at 200 rpm on a rocking platform. After 2 h, samples were centrifuged at 12000 x g for 20 min at 4°C. Supernatants were transferred to fresh tubes and kept on ice and pellets were discarded. The Bradford assay was used to quantify the protein concentration in samples. Several concentrations of bovine serum albumin (Thermo Scientific Pierce, Cat. No. PI-23209) were used to generate a standard curve, which was used to calculate protein concentrations in the lung samples. Protein Assay Dye Reagent Concentrate (Bio-Rad, Cat. No. 500-0006) was used according to the manufacturer’s instructions. Absorbance values were read at 595 nm.

### Enrichment of membrane proteins from lipopolysaccharide (LPS)-stimulated macrophages

LPS-stimulated macrophages (RAW264.7 cells and BMDM from TM mice) were collected and resuspended in 3 mL membrane lysis buffer (0.5 M EDTA, 100 mM phenylmethanesulfonylfluoride (PMSF), and 100 mM dithiothreitol in water). Samples were placed in an ice bath and cells were lysed with a probe sonicator (Cole Parmer, Illinois, USA) using 5 cycles of 10 pulses (2 s per pulse) at 40-50 A and maximum output of 8 W and checked for lysis by trypan blue exclusion. Samples were centrifuged at 700 x g for 10 min at 4°C and supernatants were transferred into new tubes and centrifuged again at 10,000 x g for 20 min at 4°C. Supernatants were collected again and centrifuged at 72,000 x g for 1 h at 4°C. The resulting pellet was resuspended in lysis buffer with 0.1% Triton-X. The protein concentration was determined using Bio-Rad RC DC Protein Assay Kit (Bio-Rad No. 500-0119) according to manufacturer’s protocol. BSA (Thermo Scientific Pierce, No. PI-23209) was used as a protein standard.

### Reverse signaling through mTNF in BMDM from TM mice

A total of 5 x 10^6^ mature BMDM generated from TM mice was resuspended in 10 mL complete DMEM10 and added to 25 cm^2^ tissue culture flasks. Cells were allowed to adhere for 2 h, prior to addition of LPS at 50 ng/mL and incubated for 6 h. To induce reverse signaling through mTNF, recombinant Crm proteins were added to individual flasks at 10 μg/mL for 3 h. Cells were then harvested for isolation of RNA, generation of cDNA and qRT-PCR analysis.

### Virus infection, animal weights and clinical scores

All mice were individually ear-tagged, weighed at the start of the experiment and then every day from day 5 p.i. Mice were anesthetized with tribromoethanol (160-240 mg/kg intraperitoneal, i.p.) and infected through the i.n. route with 25 PFU of ECTV in 30 µL PBS. The virus inoculum used to infect mice in each experiment was back-titrated and was usually found within a range of 20-60 PFU. The 50% lethal dose (LD_50_) of ECTV^WT^ for C57BL/6 WT mice infected i.n. is approximately 200 PFU. All animals were clinically assessed everyday beginning from day 1 p.i. and clinical scores recorded on a scale from 0 to 3 for each criterion, with a maximum score of 15 using a scoring system (described in Table S1) with 5 clinical parameters. For ethical reasons, mice that were severely moribund and/or lost ≥ 25% of the original body weight were euthanized and considered dead the following day.

### Treatment of mice with mAb

In some experiments, mice were administered i.p. with neutralizing/blocking mAb against IFN-γ (R4.6A2, rat IgG_1_, cat. no. BE0054), IL-6 (MP5-20F3, rat IgG_1,_ cat. no. BE0046), IL-10R (1B1.3A, rat IgG_1_, cat. no. BE0050), TNF (XT3.11, rat IgG_1_, cat. no. BE0058) or isotype control mAb (HRPN, rat IgG_1_, cat. no. BE0088) at 500 μg beginning at day 7 and every 2 days thereafter until euthanasia. Purified mAb were purchased from BioXcell (New Hampshire, USA).

### Histology and microscopic assessment of lung pathology

The left lobe of lung tissue was fixed in 10% neutral-buffered formaldehyde at room temperature for 24 h, embedded in paraffin, cut in 6 μm sagittal sections and stained with hematoxylin and eosin (H&E) for analysis with a bright field microscope (Olympus IX 71). A semi-quantitative scoring system for lung pathology was developed (described in Table S2) with 6 pathologic parameters. Individual slides were scored in a single-blinded fashion on a scale from 0 to 4 for each criterion, with a maximum score of 24.

### Flow cytometric analysis of lung leukocytes

Individual lung tissue samples were cut into small pieces and digested in EMEM containing 4 µg DNAse I, grade II and 4mg collagenase A (Roche Applied Science) and placed in a 37°C water bath for 1 h. Undigested tissue was further processed by passing through a 19-G needle attached to a syringe and then filtered through a 70 μm nylon mesh. Cells were washed with Hank’s balanced salt solution and red blood cells lysed with red cell lysis buffer (150 mM NH4Cl, 10 mM KHCO3, 0.1 mM Na2EDTA, pH 7.4) for 1 min. Cells were next washed and resuspended in PBS containing 2% FCS. FcγII/III receptors (FcγR) on cells were first blocked with anti-FcγR mAb (clone 2.4G2) for 30 min, the cells washed and stained with CD45.2-FITC (to identify leukocytes) and cell subset specific fluorochrome-conjugated mAb used in this study were obtained from BD Biosciences and Biolegend. Data were acquired on a BD LSR Fortessa flow cytometer with BD FACS Diva software and analyzed using FlowJo Software version 9.5 (Tree Star, Inc).

### Determination of cytokine protein levels

The levels of cytokines and chemokines in the lung homogenates were measured using capture ELISA kits (Biolegend), performed according to the manufacturer’s protocol. Optical density was measured at 450 nm with SOFTmax Pro software (Molecular Devices Corp).

### RNA extraction

Lung tissue from individual mice were homogenized in 1 mL of TRIsure reagent with TissueLyser II (Qiagen No. 853000) and as per the manufacturer’s instructions and samples were centrifuged for 15 min at 12,000 x g at 4°C to remove cellular debris. In the final step, RNA pellets were air-dried and resuspended in 30-50 µL RNAse-free water and incubated for 10 min at 60°C to ensure they were dissolved. RNA concentration was determined using a Nanodrop ND-1000 spectrophotometer. Samples were stored at −80°C until cDNA synthesis.

### cDNA generation

Tetro cDNA Synthesis Kit (Bioline, Cat. No. BIO-65043) was used to synthesize single-stranded cDNA from 1µg RNA in 12 µL RNAse-free water following the manufacturer’s instructions. Briefly, the priming premix was prepared on ice by mixing Oligo (dT)18 primer mix to a final concentration of 0.5 µg/µL, 10mM dNTP mix, 5x RT Buffer, 10 units/µL RiboSafe RNase Inhibitor and Tetro Reverse Transcriptase (200 unit/μL). To each sample, 8 µL of the mastermix was added and incubated at 45°C for 30 min. The reaction was terminated by incubating samples at 85°C for 5 min and then chilled on ice. Samples were diluted with 180 µL RNAse-free water and stored at −20°C until further use.

### Quantitative reverse transcription real-time polymerase chain reaction (qRT-PCR)

For each gene of interest, a real-time PCR reaction mixture was prepared containing 10 µL of SsoAdvanced™ SYBR® Green Supermix (Bio-Rad. Cat. No. 172-5265), 1 µL each of 5 µM gene-specific forward and reverse primer pair, and 6 µL of RNAse-free water. Volumes of 2 µL of the appropriate cDNA template and 18 µL of the reaction mixture were added to the wells of iQ 96-well semi-skirted PCR Plate (Bio-Rad. Cat. No. 223-9441). Amplification was achieved using Bio-Rad iQ5 Real-time PCR detection System Relative changes in gene expression was determined using delta/delta cycle threshold (2^-ΔΔCT^) analysis method (Livak and Schmittgen, 2001). Ubiquitin C (UBC) was used as the internal control. Results are reported as the fold difference relative to a calibrator cDNA (naïve mice) prepared in parallel with the experimental cDNAs.

### Viral TNFR expression and purification

Recombinant vTNFRs from ECTV (CrmD), CPXV (CrmB, CrmC and CrmD) and VARV (CrmB) were expressed in the baculovirus system, fused to a C-terminal V5-6xHis tag as described (Pontejo JBC 2015). Proteins were purified in metal chelate affinity columns from supernatants of infected Hi5 cell cultures (Pontejo JBC 2015). Permission from the World Health Organization was granted to hold VARV DNA encoding CrmB, and its manipulation was performed in accordance with the established rules.

### Statistical analysis

Statistical analyses of experimental data, as indicated in the legend to each figure, were performed using appropriate tests to compare results using GraphPad Prism 8 (GraphPad Software, Inc.). A p value of <0.05 was taken to be significant, *, p ≤ 0.05; **, p ≤ 0.01; ***, p ≤ 0.001; ****, p ≤ 0.0001.

## Supplemental Information

See separate files.

