## Supplementary material for "Poxvirus-encoded TNF receptor homolog dampens inflammation and protects the host from uncontrolled lung pathology and death during respiratory infection": Suppplementary Figures

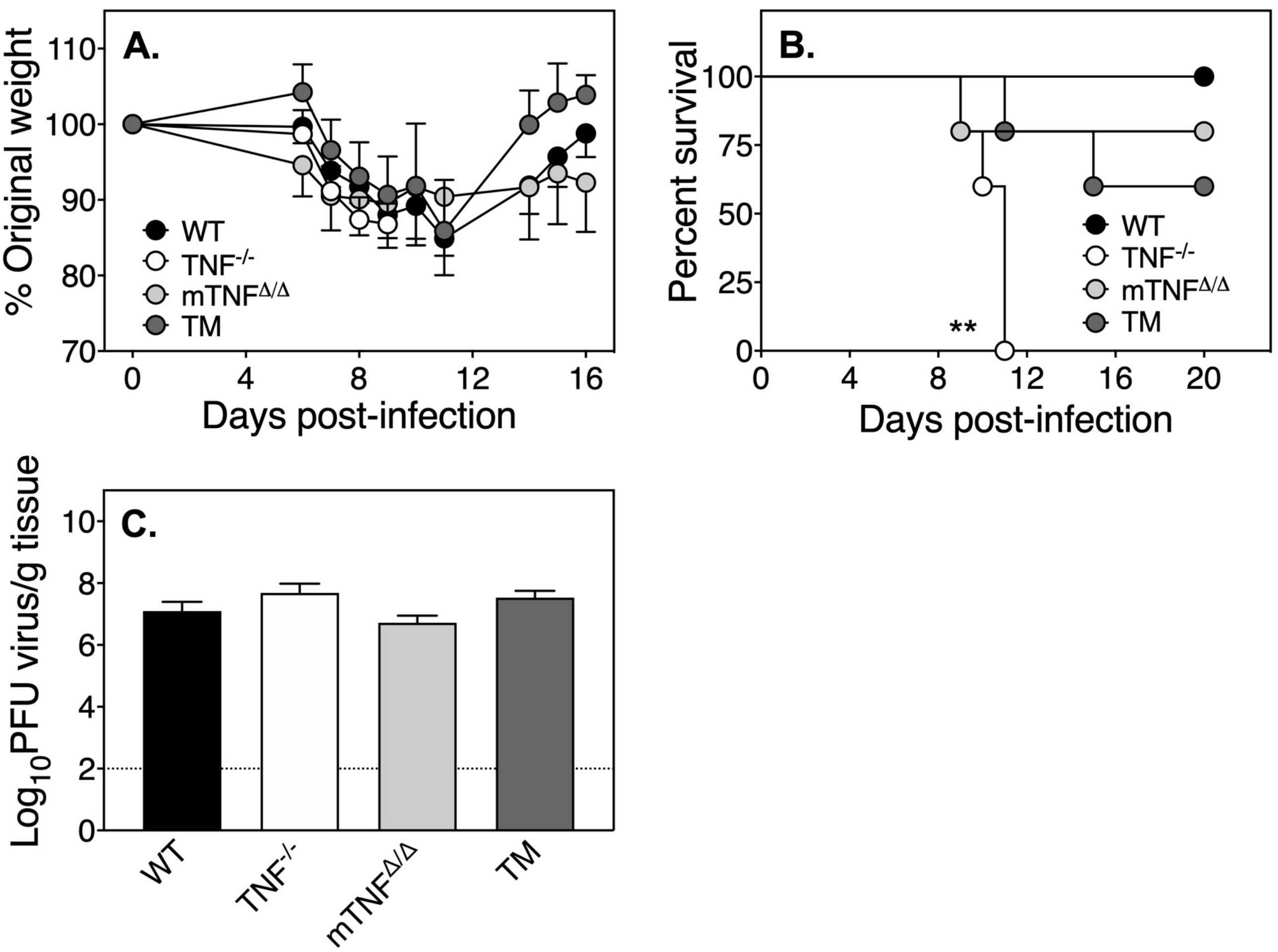


**Figure S1. Comparable resistance of mTNF$^{\Delta/\Delta}$ and TM mice to ECTV infection.** Separate groups of WT, TNF$^{-/-}$, mTNF$^{\Delta/\Delta}$ and TM mice (5 mice/group) were infected i.n. with 25 PFU ECTV$^{WT}$. One set of animals were observed for 20 days and a second group was euthanized at day 7 p.i. to measure lung viral load. Weight loss (A), survival (B), and viral load (C). Data in A are expressed as means ± SEM. For B, statistical analysis was done using the Log-rank (Mantel-Cox) test. **, $p < 0.01$ in survival proportions of WT, mTNF$^{\Delta/\Delta}$ and TM mice compared with TNF$^{-/-}$ mice. For C, data were log transformed and expressed as means ± SEM. Statistical analysis was undertaken using ordinary one-way ANOVA followed by Fisher's least significant difference tests.

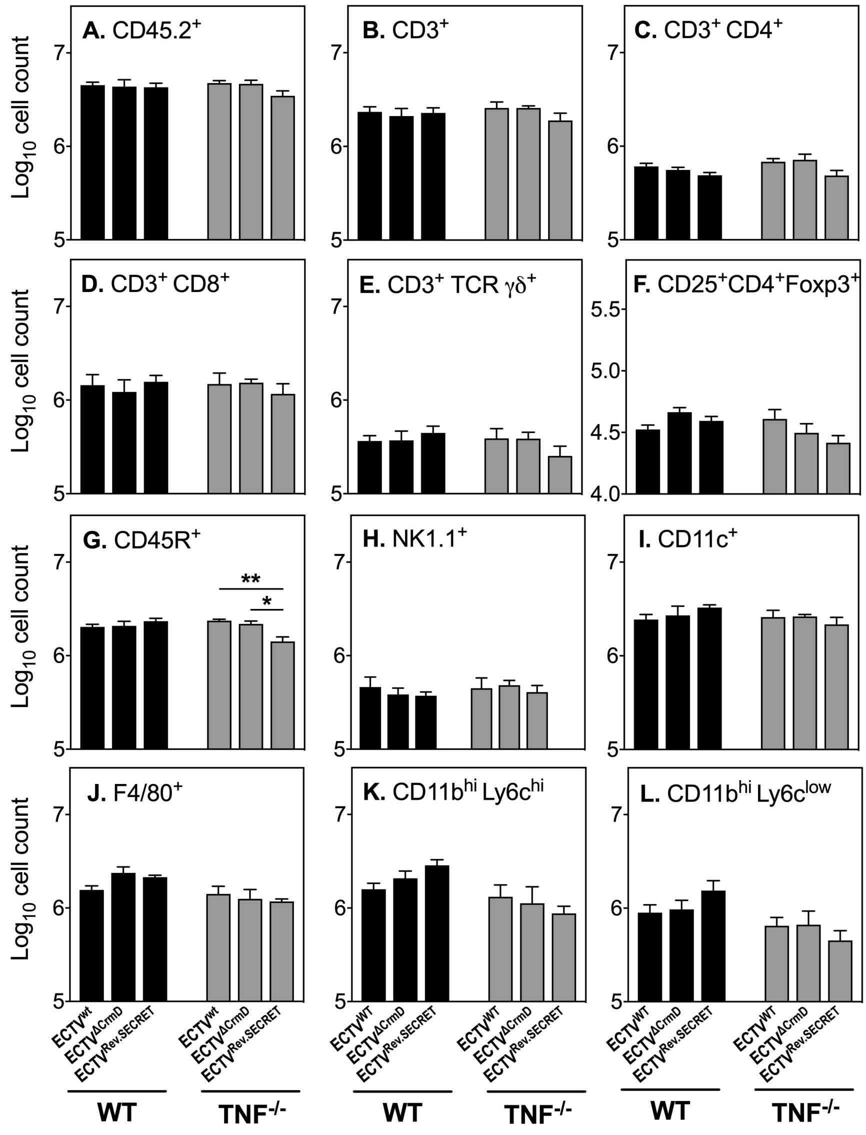


**Figure S2. Comparable leukocyte numbers and subsets in WT and TNF$^{-/-}$ at 8 days p.i. with WT or mutant viruses.** WT and TNF$^{-/-}$ mice (5 mice/group) were infected i.n. with 25 PFU ECTV$^{WT}$, ECTV$^{\Delta CrmD}$ or ECTV$^{Rev.SECRET}$. At day 8 p.i., single cell suspensions of digested lungs were stained with fluorochrome-conjugated antibodies and analyzed by flow cytometry. Shown are numbers of (A) leukocytes (CD45.2$^{+}$), (B) T cells (CD3$^{+}$), (C, D, E) T cell subsets (CD3$^{+}$CD4$^{+}$, CD3$^{+}$CD8$^{+}$ or CD3$^{+}$TCRγδ$^{+}$), (F) regulatory T cells (CD25$^{+}$CD4$^{+}$Foxp3$^{+}$), (G) B cells (CD45R$^{+}$), (H) NK cells (NK1.1$^{+}$), (I) dendritic cells (CD11c$^{+}$), (J) macrophages (F4/80$^{+}$), (K) inflammatory monocytes (CD11b$^{hi}$Ly6c$^{hi}$) and (L) neutrophils (CD45$^{+}$ CD11b$^{hi}$ Ly6c$^{low}$). Data have been log transformed and expressed as means ± SEM. Statistical significance was obtained using the two-way ANOVA followed by Tukey's multiple comparisons tests, where *, $p < 0.05$; **, $p < 0.01$; Data shown are from one of 2 separate experiments with comparable results.

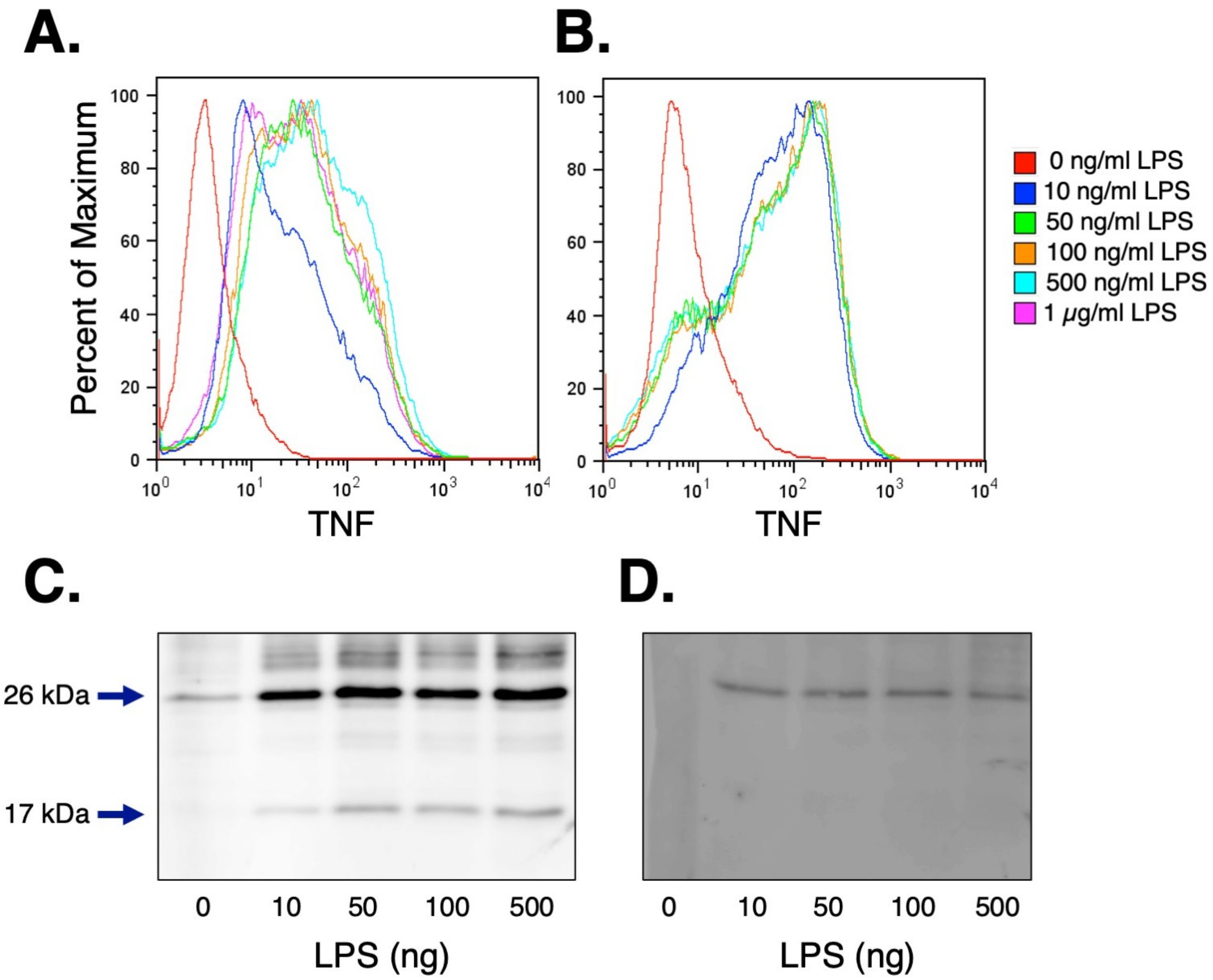


**Figure S3. Detection of mTNF on surface of macrophages stimulated with LPS.** Murine RAW264.7 cells (A) and primary BMDM from TM mice (B) were stimulated with different concentrations of LPS for 6 h. Expression of TNF on cell surface was determined by flow cytometric analysis using a fluorochrome-conjugated rat anti-mouse TNF monoclonal antibody, MP6-XT22. (C) Membrane proteins from (C) LPS-stimulated RAW264.7 cells and (D) TM BMDM were enriched and 50 μg of protein from each sample was separated on a 12% SDS-PAGE gel and transferred onto a nitrocellulose membrane. TNF was detected by Western blot analysis using an Armenian hamster anti-mouse monoclonal antibody, TN3.19-12. RAW264.7 cells produced the 26 kDa mTNF and 17 kDa sTNF while the BMDM from TM mice only produced the 26 kDa mTNF.

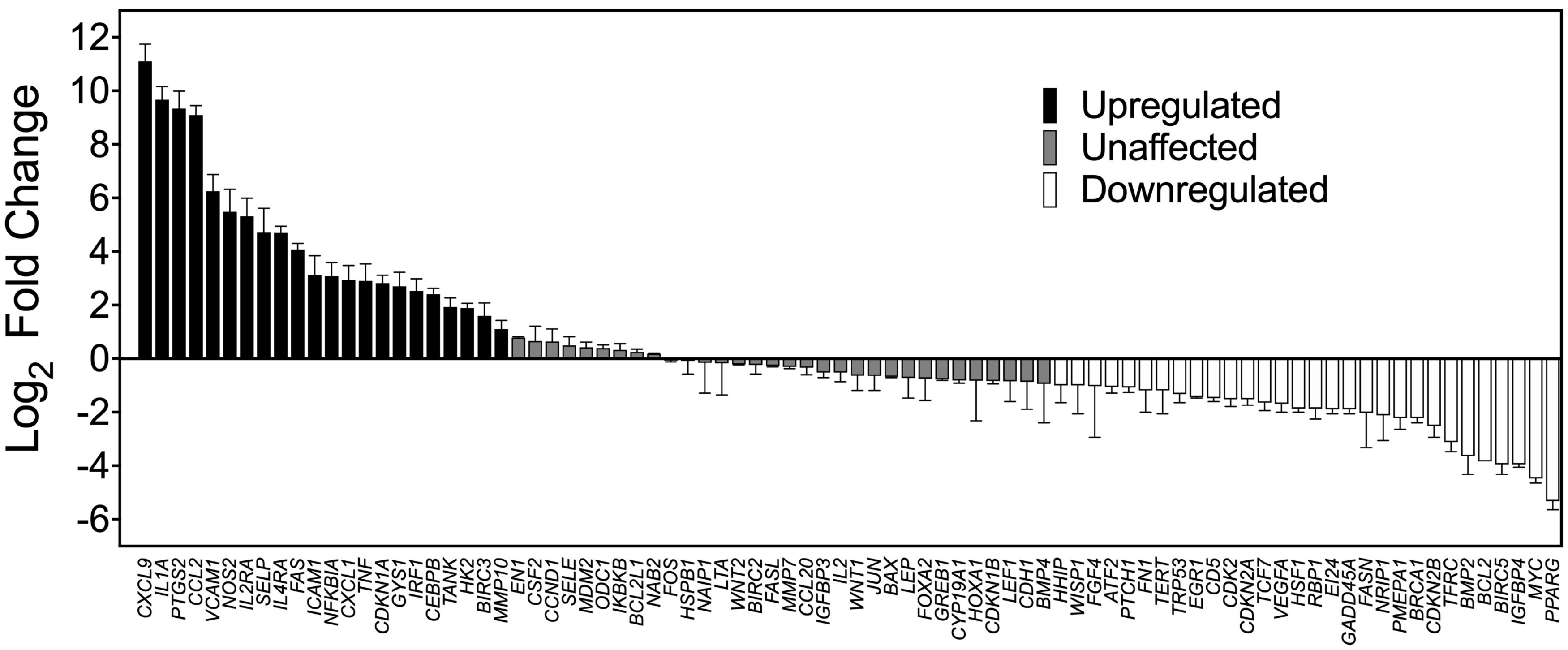


**Figure S4. Changes in gene expression profiles following LPS stimulation in BMDM from TM mice.** BMDM from TM mice were stimulated with 50 ng/mL LPS for 6 h. Expression of mRNA transcripts for cytokines was determined by RT-PCR using the SA Biosciences Pathway Finder PCR array, No. PAMM-0014. Data are combined results from two separate experiments and are expressed as means ± SEM. Five different housekeeping genes, namely, *GUSB*, *HPRT*, *HSP90AB1, GAPDH* and *ACTB* were used in the kit. The relative differences in gene expression were detected by using the delta/delta cycle threshold analysis ($2^{-\Delta\Delta CT}$) (Livak and Schmittgen, 2001). Results are expressed as fold-change compared to the unstimulated control. Reverse transcription controls to test the efficiency of reverse transcription, positive PCR controls to test the efficiency of PCR and a genomic DNA control to test DNA contamination were also included in the RT-PCR array.

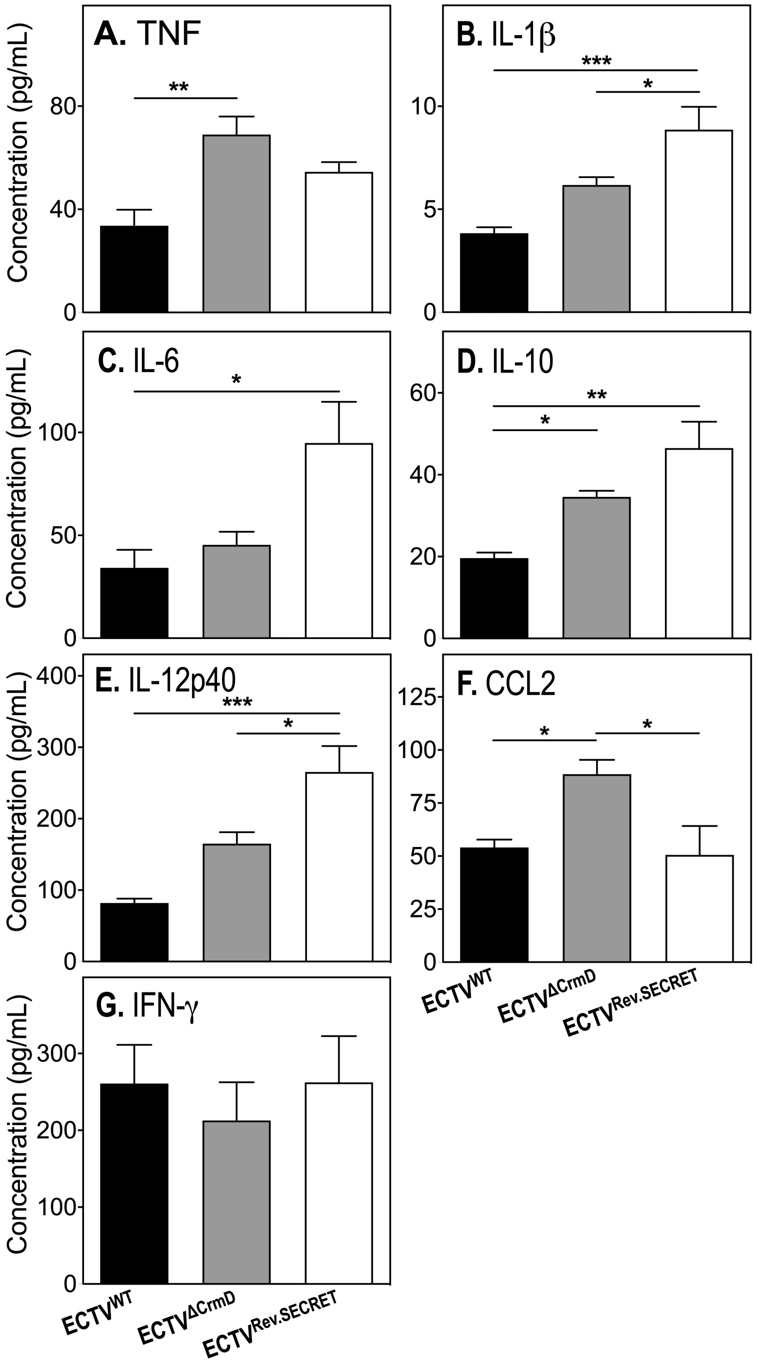


**Figure S5. Dysregulated cytokine and chemokine responses in WT mice infected with CrmD mutant ECTV.** Groups of WT mice (5 mice/group) were infected with 25 PFU i.n. with $ECTV^{WT}$, $ECTV^{\Delta CrmD}$ or $ECTV^{Rev.SECRET}$. Mice were sacrificed at day 8 p.i., lungs were collected and homogenized. Levels of (A) TNF, (B) IL-1β, (C) IL-6, (D) IL-10, (E) IL-12p40, (F) CCL2, and (G) IFN-γ in lung homogenates were measured by ELISA. Data are expressed as means ± SEM. Statistical significances were obtained by one-way ANOVA followed by Tukey's multiple comparisons tests. *, $p < 0.05$; **$p < 0.01$.
