## Supplementary Tables for "Poxvirus-encoded TNF receptor homolog dampens inflammation and protects the host from uncontrolled lung pathology and death during respiratory infection"

**Table S1**. Scoring system for clinical presentation of mice infected with ECTV

| | |
|---|---|
| I. | Hair Coat |
| | 0 – groomed and shiny<br>1 – groomed but not shiny<br>2 – rough<br>3 – unkempt ("scruffy") |
| II. | Posture |
| | 0 – back is straight<br>1 – hunched but with spontaneous straightening of the back<br>2 – hunched but straightens only upon stimulation<br>3 – hunched even when stimulated |
| III. | Breathing |
| | 0 – normal<br>1 – intermittent rapid breathing<br>2 – rapid, shallow breathing<br>3 – laboured breathing; (+) abdominal retractions |
| IV. | Lacrimation and Nasal Discharge |
| | 0 – none<br>1 – minimal lacrimation or nasal discharge<br>2 – moderate lacrimation and nasal discharge<br>3 – excessive lacrimation/catarrh; (+) blockade of either nares |
| V. | Activity/Movement/Behaviour |
| | 0 – active, spontaneous movement<br>1 – inactive, movement upon stimulation<br>2 – huddled; inactive, reluctant to move<br>3 – moribund; does not go back to prone position when placed on a supine position |

**Table S2**. Histopathological assessment of lung sections from ECTV-infected animals

| | |
|---|---|
| I. | Parenchymal/Intra-alveolar Edema |
| | 0 – None |
| | 1 – Accumulation of fluid in <25% of pulmonary parenchyma |
| | 2 – Accumulation of fluid in 25-50% of pulmonary parenchyma |
| | 3 – Accumulation of fluid in 50-75% of pulmonary parenchyma |
| | 4 – Accumulation of fluid in >75% of pulmonary parenchyma |
| II. | Perivascular Edema |
| | 0 – None |
| | 1 – Mild accumulation of fluid in few perivascular spaces |
| | 2 – Mild to moderate accumulation of fluid in some perivascular spaces |
| | 3 – Moderate to severe accumulation of fluid in most of the perivascular spaces |
| | 4 – Moderate to severe accumulation of fluid in all of the perivascular spaces |
| III. | Degree of Bronchial Epithelium Necrosis |
| | 0 – None |
| | 1 – < 25% epithelial necrosis of few bronchioles |
| | 2 – < 25% epithelial necrosis of most bronchioles |
| | 3 – 25-50% epithelial necrosis of most bronchioles |
| | 4 – > 50% epithelial necrosis of most bronchioles |
| IV. | Parenchymal Inflammatory Infiltrates |
| | 0 – Few infiltrates in few areas |
| | 1 – Only few infiltrates in separate areas/foci |
| | 2 – Many scattered infiltrates in few areas/foci |
| | 3 – Confluent infiltrates in few areas/foci |
| | 4 – Diffuse inflammatory infiltrates |
| V. | Perivascular Inflammatory Infiltrates |
| | 0 – Few infiltrates in few spaces |
| | 1 – Only few infiltrates in most spaces |
| | 2 – Many scattered infiltrates in few perivascular spaces |
| | 3 – Confluent infiltrates in few perivascular spaces |
| | 4 – Infiltrates occupying most of the areas of the spaces |
| VI. | Alveolar Septal Wall Damage |
| | 0 – None |
| | 1 – Damage in <25% of the alveolar septal walls |
| | 2 – Damage in 25-50% of the alveolar septal walls |
| | 3 – Damage in 50-75% of the alveolar septal walls |
| | 4 – Damage in >75% of the alveolar septal walls |

**Table S3**. Gene List for Mouse Signal Transduction PathwayFinder PCR Array from NCBI GenBank (Benson et al. 2011)

| GeneBank ID | Symbol | Description |
|---|---|---|
| NM_009715 | *ATF2* | Activating transcription factor 2 |
| NM_007527 | *BAX* | Bcl2-associated X protein |
| NM_009741 | *BCL2* | B-cell leukemia/lymphoma 2 |
| NM_009743 | *BCL2L1* | Bcl2-like 1 |
| NM_008670 | *NAIP1* | NLR family, apoptosis inhibitory protein 1 |
| NM_007465 | *BIRC2* | Baculoviral IAP repeat-containing 2 |
| NM_007464 | *BIRC3* | Baculoviral IAP repeat-containing 3 |
| NM_009689 | *BIRC5* | Baculoviral IAP repeat-containing 5 |
| NM_007553 | *BMP2* | Bone morphogenetic protein 2 |
| NM_007554 | *BMP4* | Bone morphogenetic protein 4 |
| NM_009764 | *BRCA1* | Breast cancer 1 |
| NM_011333 | *CCL2* | Chemokine (C-C motif) ligand 2 |
| NM_016960 | *CCL20* | Chemokine (C-C motif) ligand 20 |
| NM_007631 | *CCND1* | Cyclin D1 |
| NM_007650 | *CD5* | CD5 antigen |
| NM_009864 | *CDH1* | Cadherin 1 |
| NM_016756 | *CDK2* | Cyclin-dependent kinase 2 |
| NM_007669 | *CDKN1A* | Cyclin-dependent kinase inhibitor 1A (P21) |
| NM_009875 | *CDKN1B* | Cyclin-dependent kinase inhibitor 1B |
| NM_009877 | *CDKN2A* | Cyclin-dependent kinase inhibitor 2A |
| NM_007670 | *CDKN2B* | Cyclin-dependent kinase inhibitor 2B (p15, inhibits CDK4) |
| NM_009883 | *CEBPB* | CCAAT/enhancer binding protein (C/EBP), beta |
| NM_009969 | *CSF2* | Colony stimulating factor 2 (granulocyte-macrophage) |
| NM_008176 | *CXCL1* | Chemokine (C-X-C motif) ligand 1 |
| NM_008599 | *CXCL9* | Chemokine (C-X-C motif) ligand 9 |
| NM_007810 | *CYP19A1* | Cytochrome P450, family 19, subfamily a, polypeptide 1 |
| NM_007913 | *EGR1* | Early growth response 1 |
| NM_007915 | *EI24* | Etoposide induced 2.4 mRNA |
| NM_010133 | *EN1* | Engrailed 1 |
| NM_007987 | *FAS* | Fas (TNF receptor superfamily member 6) |
| NM_010177 | *FASL* | Fas ligand (TNF superfamily, member 6) |
| NM_007988 | *FASN* | Fatty acid synthase |
| NM_010202 | *FGF4* | Fibroblast growth factor 4 |
| NM_010233 | *FN1* | Fibronectin 1 |
| NM_010234 | *FOS* | FBJ osteosarcoma oncogene |
| NM_010446 | *FOXA2* | Forkhead box A2 |
| NM_007836 | *GADD45A* | Growth arrest and DNA-damage-inducible 45 alpha |
| NM_015764 | *GREB1* | Gene regulated by estrogen in breast cancer protein |
| NM_030678 | *GYS1* | Glycogen synthase 1, muscle |
| NM_020259 | *HHIP* | Hedgehog-interacting protein |
| NM_013820 | *HK2* | Hexokinase 2 |
| NM_010449 | *HOXA1* | Homeo box A1 |
| NM_008296 | *HSF1* | Heat shock factor 1 |
| NM_013560 | *HSPB1* | Heat shock protein 1 |
| NM_010493 | *ICAM1* | Intercellular adhesion molecule 1 |
| NM_008343 | *IGFBP3* | Insulin-like growth factor binding protein 3 |
| NM_010517 | *IGFBP4* | Insulin-like growth factor binding protein 4 |
| NM_010546 | *IKBKB* | Inhibitor of kappaB kinase beta |
| NM_010554 | *IL1A* | Interleukin 1 alpha |
| NM_008366 | *IL2* | Interleukin 2 |
| NM_008367 | *IL2RA* | Interleukin 2 receptor, alpha chain |
| NM_001008700 | *IL4RA* | Interleukin 4 receptor, alpha |
| NM_008390 | *IRF1* | Interferon regulatory factor 1 |
| NM_010591 | *JUN* | Jun oncogene |
| NM_010703 | *LEF1* | Lymphoid enhancer binding factor 1 |
| NM_008493 | *LEP* | Leptin |

| | | |
|---|---|---|
| NM_010735 | *LTA* | Lymphotoxin A |
| NM_010786 | *MDM2* | Transformed mouse 3T3 cell double minute 2 |
| NM_019471 | *MMP10* | Matrix metallopeptidase 10 |
| NM_010810 | *MMP7* | Matrix metallopeptidase 7 |
| NM_010849 | *MYC* | Myelocytomatosis oncogene |
| NM_008668 | *NAB2* | Ngfi-A binding protein 2 |
| NM_010907 | *NFKBIA* | Nuclear factor of kappa light polypeptide gene enhancer in B-cells inhibitor, alpha |
| NM_010927 | *NOS2* | Nitric oxide synthase 2, inducible |
| NM_173440 | *NRIP1* | Nuclear receptor interacting protein 1 |
| NM_013614 | *ODC1* | Ornithine decarboxylase, structural 1 |
| NM_011146 | *PPARG* | Peroxisome proliferator activated receptor gamma |
| NM_008957 | *PTCH1* | Patched homolog 1 |
| NM_011198 | *PTGS2* | Prostaglandin-endoperoxide synthase 2 |
| NM_011254 | *RBP1* | Retinol binding protein 1, cellular |
| NM_011345 | *SELE* | Selectin, endothelial cell |
| NM_011347 | *SELP* | Selectin, platelet |
| NM_011529 | *TANK* | TRAF family member-associated Nf-kappa B activator |
| NM_009331 | *TCF7* | Transcription factor 7, T-cell specific |
| NM_009354 | *TERT* | Telomerase reverse transcriptase |
| NM_011638 | *TFRC* | Transferrin receptor |
| NM_022995 | *PMEPA1* | Prostate transmembrane protein, androgen induced 1 |
| NM_013693 | *TNF* | Tumor necrosis factor |
| NM_011640 | *TRP53* | Transformation related protein 53 |
| NM_011693 | *VCAM1* | Vascular cell adhesion molecule 1 |
| NM_009505 | *VEGFA* | Vascular endothelial growth factor A |
| NM_018865 | *WISP1* | WNT1 inducible signaling pathway protein 1 |
| NM_021279 | *WNT1* | Wingless-related MMTV integration site 1 |
| NM_023653 | *WNT2* | Wingless-related MMTV integration site 2 |

**Table S4.** Genes downregulated by LPS stimulation and modulated following reverse signaling with vTNFRs

| Genes | Pathway | Fold change relative to unstimulated cells | | | | |
|---|---|---|---|---|---|---|
| | | ECTV crmD | CPXV crmB | CPXV crmC | CPXV crmD | VARV crmB |
| *HSF1* | Stress | -3.47 | 1.69 | 1.36 | 1.97 | -0.79 |
| *HHIP* | Hedgehog | -0.15 | -1.16 | -1.94 | 2.45 | 1 |
| *BIRC5* | Wnt | 1.06 | 1.12 | -0.83 | 3.61 | -1.24 |
| *TERT* | NF-κB | 1.19 | -2.95 | -0.95 | -0.30 | 1.48 |
| *EI24* | p53 | 1.26 | 1.3 | 1.24 | 2.73 | -2.24 |
| *EGF1* | CREB | 1.31 | -3.57 | -2.72 | -2.12 | -0.40 |
| *TCF7* | Wnt | 1.31 | -0.03 | -0.83 | 3.61 | 1 |
| *FASN* | Metabolism | 1.33 | -4.64 | -2.95 | -0.87 | -1.43 |
| *CDK2* | Androgen | 1.44 | 1.55 | 1.38 | -2.87 | -0.64 |
| *PPARG* | Wnt | 1.49 | 1.07 | 1.07 | -2.72 | 1.47 |
| *IGFBP4* | Estrogen | 1.52 | 1.28 | 1.35 | 2.28 | -0.95 |
| *WISP1* | Wnt | 1.59 | 2.11 | -0.99 | 3.46 | -3.38 |
| *BRCA1* | Estrogen | 1.63 | 1.2 | -2.18 | 1.4 | -1.03 |
| *PMEPA1* | Androgen | 1.63 | 2.48 | 1.29 | 3.61 | 1 |
| *BMP2* | Hedgehog | 1.91 | -0.03 | -0.83 | 3.61 | 1 |
| *FN1* | PIA3K/AKT | 2.03 | 1.06 | -4.79 | -1.33 | -3.38 |
| *PTCH1* | Hedgehog | 2.13 | 2.01 | 1.3 | -0.64 | 3.01 |
| *NRIP1* | Estrogen | 2.19 | -1.83 | -1.24 | -2.64 | -1.29 |
| *CD5* | NFAT | 2.89 | -0.54 | -0.83 | 1.3 | -1.11 |
| *BCL2* | PLC | 2.91 | 1.3 | 1.86 | 2.6 | -2.18 |
| *FGF4* | Wnt | 3.25 | -0.03 | -0.83 | 3.61 | 1 |
| *CDKN2B* | TGF-β | 3.51 | -1.24 | 1 | 1.07 | 1.39 |
| *RBP1* | Retinoic Acid | 3.92 | -0.03 | -0.83 | 3.61 | 1 |
| *MYC* | Stress | 3.95 | 2.08 | 2.54 | -0.44 | -0.37 |

(orange) ≥2-fold decrease in gene expression
(green) ≥2-fold increase in gene expression
